# A Parasite Odyssey: An RNA virus concealed in *Toxoplasma gondii*

**DOI:** 10.1101/2023.09.17.558162

**Authors:** Purav Gupta, Aiden Hiller, Jawad Chowdhury, Declan Lim, Dillon Yee Lim, Jeroen P.J. Saeij, Artem Babaian, Felipe Rodriguez, Luke Pereira, Alex Morales

**Author notes:** Authors contributed equally, order was determined by Starfox64 tournament.

## Abstract

We are entering a “Platinum Age of Virus Discovery”, an era marked by exponential growth in the discovery of virus biodiversity, and driven by advances in metagenomics and computational analysis. In the ecosystem of a human (or any animal) there are more species of viruses than simply those directly infecting the animal cells. Viruses can infect all organisms constituting the microbiome, including bacteria, fungi, and unicellular parasites. Thus the complexity of possible interactions between host, microbe, and viruses is unfathomable. To understand this interaction network we must employ computationally-assisted virology as a means of analyzing and interpreting the millions of available samples to make inferences about the ways in which viruses may intersect human health.

From a computational viral screen of human neuronal datasets, we identified a novel narnavirus *Apocryptovirus odysseus* (Ao) which likely infects the neurotropic parasite *Toxoplasma gondii*. Previously, several parasitic protozoan viruses (PPVs) have been mechanistically established as triggers of host innate responses, and here we present *in silico* evidence that Ao is a plausible pro-inflammatory factor in human and mouse cells infected by *T. gondii*. *T. gondii* infects billions of people worldwide, yet the prognosis of toxoplasmosis disease is highly variable, and PPVs like Ao could function as a hitherto undescribed hypervirulence factor. In a broader screen of over 7.6 million samples, we explored phylogenetically-proximal viruses to Ao and discovered 19 *Apocryptovirus* species, all found in libraries annotated as vertebrate transcriptome or metatranscriptomes. While the Narnavirus samples making up this genus-like clade are derived from sheep, goat, bat, rabbit, chicken, and pigeon samples, the presence of virus is strongly predictive of parasitic (*Apicomplexa*) nucleic acid co-occurrence, supporting that these are a clade of parasite-infecting viruses.

This is a computational proof-of-concept study in which we rapidly analyze millions of datasets from which we distilled a mechanistically, ecologically, and phylogenetically refined hypothesis. We predict this highly diverged Ao RNA virus is biologically a *T. gondii* infection, and that Ao, and other viruses like it, will modulate this disease which afflicts billions worldwide.

## Introduction

RNA virus discovery is undergoing a revolutionary expansion in the characterization of virus diversity (Shi et al., 2016, 2018; Wolf et al., 2020; Edgar et al., 2022; Neri et al., 2022; Zayed et al., 2022; Charon et al., 2022; Olendraite et al., 2023; Hou et al., 2023; Lee et al., 2023; Forgia et al., 2023; Zheludev et al., 2024). Of the predicted 10^8^ to 10^12^ virus species on Earth (Koonin et al., 2020), *≈* 300, 000 mammalian viruses are thought to have human-infecting potential (Anthony et al., 2013), of which we know *≈* 160 RNA viruses (Woolhouse and Adair, 2013). This bulk of unknown wildlife-originating (zoonotic) viruses are expected to cause the majority of emerging infectious diseases in humans (Jones et al., 2008), with precedent set by the 1918 Spanish influenza, AIDS, SARS, Ebola, and more recently COVID-19. This establishes the real and continued threat that viral zoonoses pose to global health.

Most established relationships between a disease and its causal RNA virus are direct and proximal to infection and are thus tractable to reductionist interrogation. Yet it is evident that confounding variables can impede linking cause and effect: virus genetic heterogeneity, chronic infections, asymptomatic carriers, prolonged latency periods, and microbiome interactions all add complexity to the link between virus and disease. An indirect, yet causal, relationship is well exemplified by Epstein-Barr virus and multiple sclerosis (EBV-MS). The EBV-MS association has long been statistically postulated, yet clear evidence for causality was only established recently, on a background of increasing awareness of the role of neuroinflammation in neurodegeneration (Bjornevik et al., 2022; Lanz et al., 2022; Soldan and Lieberman, 2023). Thus, statistical and computational inferences, while not sufficient to formalize causation, do allow for a radical simplification of the space of plausible hypotheses, and thereby accelerates the time to discovery. We opine that in addition to virus discovery, virus effect inference, pathological and ecological, should be a primary objective of the emerging field of computational virology.

### Asking an old question in a new way

The volume of freely available sequencing data in the Sequence Read Archive (SRA) has grown explosively for over a decade (Katz et al., 2021). Currently, there are in excess of 52.96 petabases (Pbp) of freely available sequencing data, capturing *>* 30 million biological samples. The emerging field of petabase-scale computational biology strives to analyze the totality of this data and enable expansive meta-analyses. The SRA-STAT project has processed over 10.8 million datasets (27.9 Pbp) using a k-mer hashing algorithm to create an index of reads matching known taxa genomes. (Katz et al., 2021). Likewise, Serratus which is aimed at uncovering known and novel RNA viruses using a translated nucleotide search for the RNA-dependent RNA polymerase (RdRp), the hallmark gene of RNA viruses, has processed over 7.5 million RNAsequencing datasets (18.97 Pbp) (Edgar et al., 2022). Our group focuses on leveraging this critical mass of freely available data to re-interrogate fundamental questions in virology using a data-driven philosophy. This approach allows us to minimize *a priori* bias and maximize the discovery of unexpected biology. Our immediate focus is to characterize highly divergent neuroinflammatory RNA viruses - these, we hypothesize, have the potential to cause or contribute to human neurological diseases of unknown aetiology.

### As simple as it gets - the Narnaviridae

One group of poorly understood yet highly divergent viruses are the *Narnaviridae*. This clade are among the simplest viruses, comprising a naked, +ve sense RNA genome (hence the derivation from ”naked RNA”) encoding an RNA-dependent RNA polymerase; the virus is thus observed as a ribonucleoprotein complex, with no true virion (Hillman and Esteban, 2011; Hillman and Cai, 2013). Members of this family of viruses are best known for their association with fungi, and indeed the first two species identified, Saccharomyces 20S RNA virus (ScNV-20S) and Saccharomyces 23S RNA virus (ScNV-23S) were discovered in the model organism *Saccharomyces cerevisiae* (Kadowaki and Halvorson, 1971; Wejksnora and Haber, 1978; Garvik and Haber, 1978). ScNV-20S and ScNV-23S infections are mostly not associated with a phenotype, (Hillman and Esteban, 2011; Hillman and Cai, 2013) although as is the case with the *S. cerevisiae* L-A virus, chronic apathogenic infections may become phenotypic in specific genetic backgrounds (Chau et al., 2023). Likewise, although *S. cerevisiae* strains harbouring high expression of a related narnavirus, N1199, display defects in sporulation, autophagy and a change in metabolite utilisation, strains with a low N1199 expression are more common and display no phenotype (Vijayraghavan et al., 2023). A comparison of virus-infected and virus-eliminated strains of *Aspergillus flavus* found that narnavirus infection is not associated with changes in colony appearance, growth rate or mycelial/hyphal morphology, despite changes in transcriptomic profile (Kuroki et al., 2023). Besides fungi, *Narnaviridae* have also been found in marine protists (Charon et al., 2021; Chiba et al., 2023), mosquitoes (Batson et al., 2021; Yang et al., 2023; Abbo et al., 2023), and other arthropods (Harvey et al., 2019; Xu et al., 2022).

### Parasitic protozoan viruses, nested invaders

The niches of the human virome extend beyond human cells; our holobiont constitutes an array of biological hosts including the bacteria, fungi, plants, and parasites. Of interest is the capacity of diverse viruses to modulate the physiology of these non-human hosts and in doing so, influence the biology of the ”macrohost” - *Homo sapiens* (Gömez-Arreaza et al., 2017; Zhao et al., 2023; Heeren et al., 2023). Perhaps unsurprisingly, bacteria and their bacteriophages dominate the human microbiome and have been the focus of the majority of metagenomic research to date (Khan Mirzaei et al., 2021; Liang and Bushman, 2021). Yet one intriguing category of human-adjacent viruses are parasitic protozoan viruses (PPVs) which infect the eukaryotic phyla *Euglenozoa* and *Apicomplexa* (Lye et al., 2016; Grybchuk et al., 2018; Charon et al., 2019; Rodrigues et al., 2022). PPVs are a functional (and diverse), rather than phylogenetic grouping and are generally poorly characterized (Gömez-Arreaza et al., 2017; Zhao et al., 2023; Heeren et al., 2023).

More than mere passengers, the presence of a PPV within a parasite and subsequent exposure of the macrohost to PPV-derived pathogen-associated molecular patterns (PAMPs) can modulate macrohost immune responses, with important implications for pathogenicity (Gömez-Arreaza et al., 2017; Zhao et al., 2023). Viral PAMPs, including viral RNA (vRNA), are sufficient to initiate an innate immune response via nucleic acid sensors (NAS), even in the context of parasitic infection of the macrohost. NAS include toll-like receptors (TLRs) which survey endosomes for dsRNA (TLR3), ssRNA (TLR7,8), or unmethylated CpG motifs in ssDNA (TLR9) (Fitzgerald and Kagan, 2020). Additionally, the three RIG-I-like receptors (RLRs) - the signalling RIG-I and MDA5, and the regulatory LGP2 - survey the cytosol for vRNA transcripts with an exposed 5’ triphosphate, or misprocessed cellular RNA (Rehwinkel and Gack, 2020). Collectively, NASs orchestrate an antiviral type I interferon (IFN) response (Lee and Ashkar, 2018). For example, the dsRNA virus Cryptosporidium parvum virus 1 (CSpV1) which infects *Cryptosporidium parvum*, activates a type I IFN inflammatory cascade in mouse and cell culture models. Interestingly, this response undermines host defences against *C. parvum*, as evidenced by the enhanced antiparasitic immunity observed in mice lacking type I IFN receptor in their intestinal epithelia (Deng et al., 2023). While the underlying mechanism for the inflammation remains elusive, PPV presence appears to be necessary for parasite pathogenicity, potentially by diverting the host’s innate immune system towards the activation of antiviral immunity and away from antiparasitic immunity. The activation of a NAS-dependent type I IFN response by a PPV is also observed with Trichomonas vaginalis virus (TVV) infecting *Trichomonas vaginalis* and Leishmania RNA virus infecting *Leishmania sp.* (Fichorova et al., 2012; Narayanasamy et al., 2022; de Carvalho et al., 2019). In both of these common human pathogens, the virus is predicted or observed to worsen the severity of parasitic disease. Conversely, PPVs, such as Giardia lamblia virus 1 (GLV1) can also impair parasitic pathogenesis. In the case of GLV1, the virus inhibits the growth of *G. lamblia* (Miller et al., 1988), highlighting the complexity of this tripartite relationship. No such relationships, however, have been observed in narnavirus or narnavirus-like viruses.

Surprisingly, there are no known viruses associated with *Toxoplasma gondii* (*T. gondii*), an apicomplexan parasite that infects approximately 30% of the global human population, with some geographic regions reaching majority seroprevalence (Pappas et al., 2009; Dubey, 2021a; Bisetegn et al., 2023). *T.gondii* has a notably broad host range of warm-blooded animals, and tropism for all animal tissues, including the brain (Pappas et al., 2009; Dubey, 2021a). While *T. gondii* infection in *Homo sapiens* is most commonly asymptomatic and self-limiting, it remains a dangerous, opportunistic infection in immunocompromised patients and pregnant women, and a leading infectious cause of blindness (Khurana and Batra, 2016; Goh et al., 2023). Furthermore, sporadic *T. gondii* strains are hypervirulent, with the ability to cause severe disease even in immunocompetent individuals (Dardé et al., 2020). Thus the full global health burden of toxoplasmosis remains unclear, especially its possible role in chronic neuroinflammation and subsequent neurodegenerative disease.

### Tell me, O Muse - a viral Odyssey

In this work, we use the SRA and Serratus to screen for potential human neuroinflammatory viruses, identifying *Apocryptovirus odysseus*, a narnavirus hidden within *Toxoplasma gondii*, and 19 additional members of the proposed genus *Apocryptovirus*, all of which likely infect apicomplexan parasites. We establish these viruses in their phylogenetic context, estimate their prevalence, and perform sequence and structure analysis of the *Apocryptovirus* RdRp. Finally, we provide computational evidence and describe a model for *Apocryptovirus odysseus*-mediated *Toxoplasma* hypervirulence.

## Results

### Identification of T. gondii associated viral sequences

Using Serratus to screen for novel human neuroinflammatory RNA viruses, we searched the BioSample database (Barrett et al., 2012) for SRA-libraries annotated with meta-data describing the sample as (1) human and (2) central nervous system (see Materials and Methods). From the 7,675,502 sequencing runs or 18.97 Pbp processed in the Serratus database, 483,173 (6.29%) runs were annotated as central nervous system samples, of which 82,454 (17.07%) were annotated as human. We further filtered to “neuron” annotated datasets containing an unknown virus (*≤*90% RdRp amino acid identity as detected by palmscan (Babaian and Edgar, 2022), which yielded a shortlist of 10 matching libraries. By contig coverage, 3 of the highest-expression libraries (SRR1205923, SRR1204654, and SRR1205190) originated from one BioProject (PRJNA241125), which we manually inspected to identify a divergent RdRp fragment (palmprint: u150420 halalDiploma). We have termed the BioProject’s associated paper “the Nĝo study”, wherein the authors infected three human cell types (neuronal stem cells, neuronal differentiated cells, and monocytes) *in vitro*, with nine different strains of the apicomplexan parasite *T. gondii* or a mock control (Nĝo et al., 2017). We assembled all 117 available Nĝo mRNAseq runs *de novo* and with palmscan identified a 3,177 nucleotide (nt) putative viral contig with a coding-complete 989 amino acid RdRp Figure 1. A BLASTp search on the translated protein revealed highly divergent RdRp homologs, the highest identity match was an unclassified *Riboviria* RdRp (date: 2023-06-18, accession: QIM73983.1, percent-identity: 54.13%, query-coverage: 40%, E-value: 9e-133).

**Figure 1.**
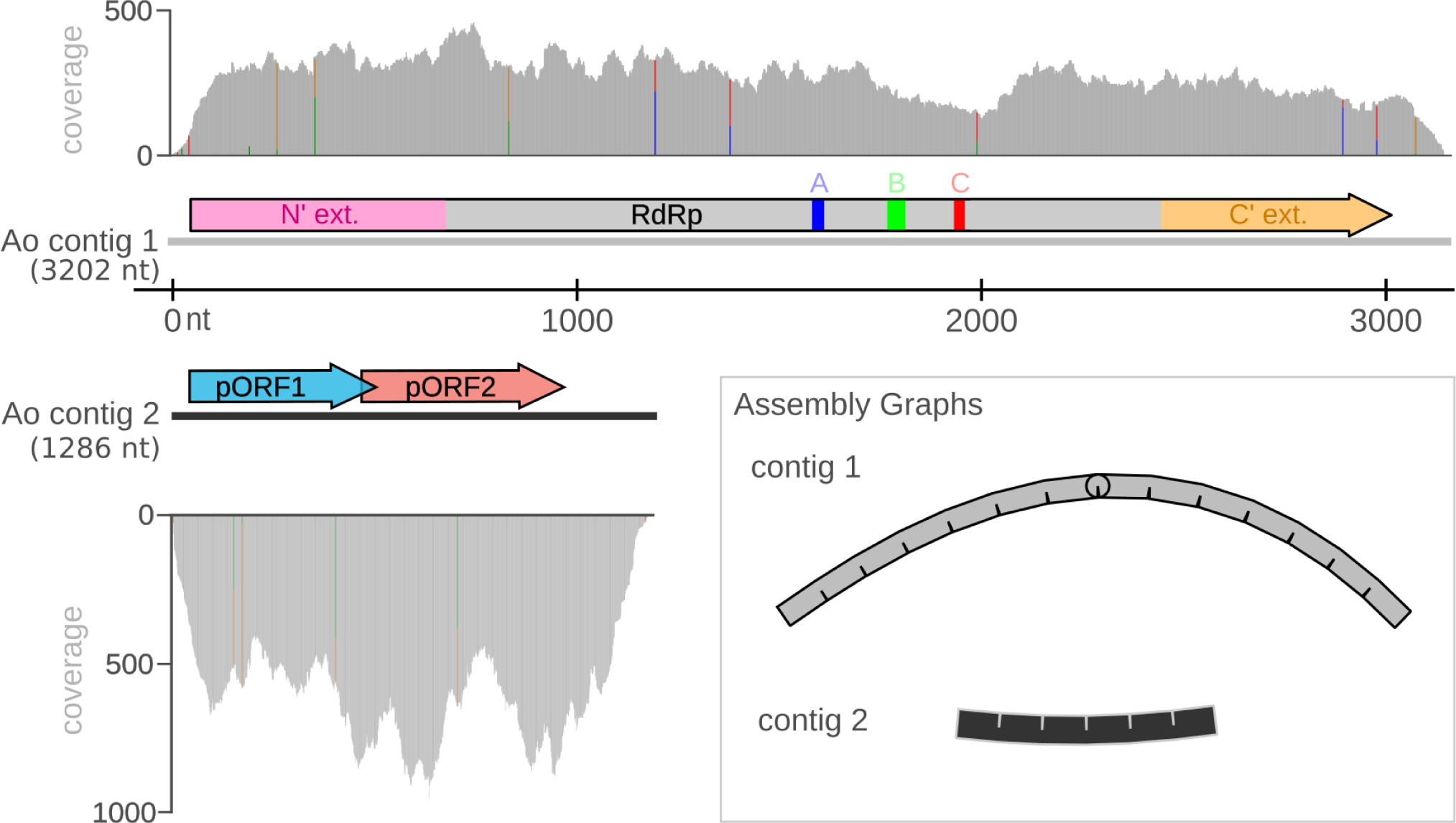
*Apocryptovirus odysseus* (Ao) genome. From (rnaSPAdes 3.15.5) assembly of the the Nĝo RNA sequencing dataset (SRR1205923), we recovered a high-coverage (123x coverage) RdRp-encoding contig, and identified a second correlated contig of likely viral origin. Contig 2 (279x coverage) contains two putative ORFs: pORF1 and pORF2 but these ORFs do not have identifiable homologs via BLASTp in the non-redundant and transcriptome shotgun assembly databases (date accessed: June 2023), TBLASTN with the nucleotide database, nor by InterProScan or HHpred (dates accessed: June 2023). Inlay, unbranched assembly graphs (Bandage 0.8.1) of both contigs confirmed a linear genome structure.

We then screened for additional viral segments or co-occurring contigs by depleting contigs mapping to the genomes of either *T. gondii* or *H. sapiens* (Materials and Methods subsection 4.3). This produced a second 1,283 nt contig whose expression correlated with the RdRp (Pearson correlation *R* = 0.95) (Figure 2C). This contig encodes four open reading frames (ORFs) with no identifiable homologs (Figure 1). Given the well-conserved RdRp sequence is already highly divergent, it is not unexpected that a viral accessory or structural gene would prove more difficult to identify based on sequence homology.

**Figure 2.**
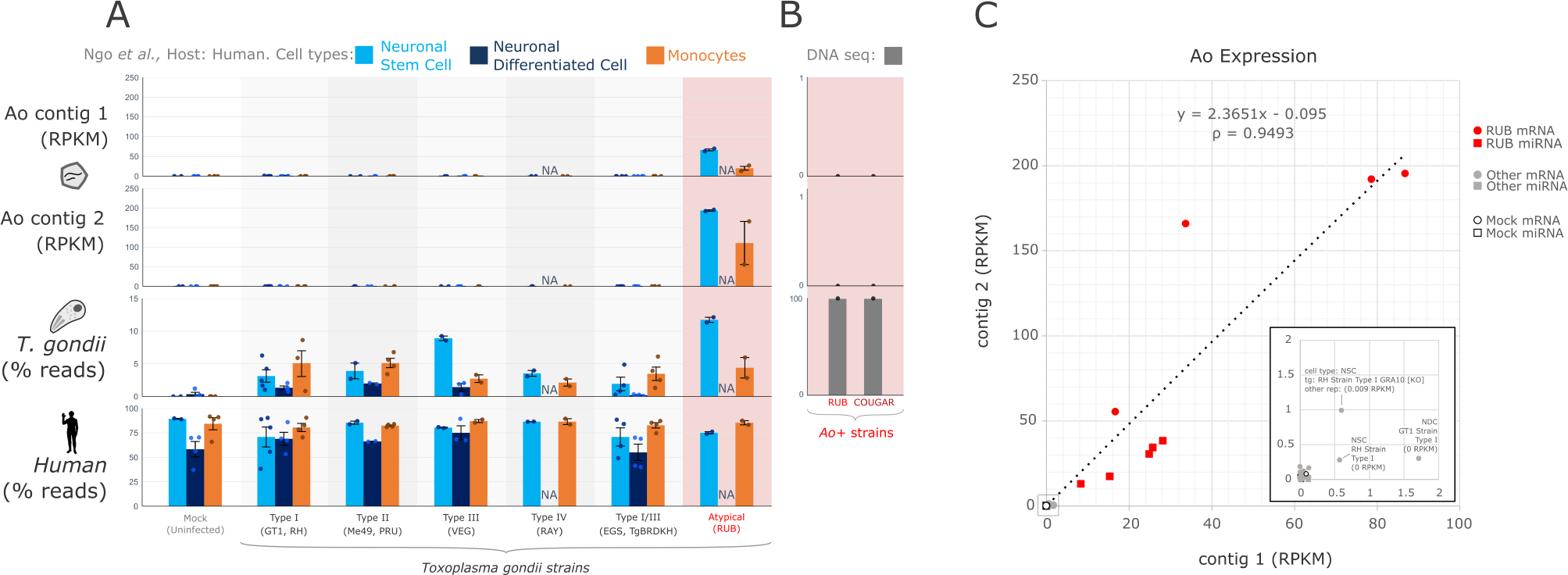
Quantification of *A. odysseus* (Ao) genome across Toxoplasma-infected human cell lines in the Ngô study. **A.** The expression (reads per million kilobase of mapped reads, RPKM) of contig 1 and contig 2 of *A. odysseus* (Ao) was quantified in three cell types (n = 37 neuronal stem cell, NSC; n = 24 neuronal differentiated cell, NDC; and n = 40 monocytes), infected by one of nine strains (n = 117 runs total) of *T. gondii*, in addition to quantifying Toxoplasma or human genome-mapped reads (percentage total) from the same datasets. Ao was found at high expression exclusively in *T. gondii* - RUB strain samples within Ngô *et al*. mRNAseq and miRNAseq (not shown). Data points and 2 SD Error Bars are shown. **B.** DNAseq quantification of Ao in four replicates of *T. gondii* - RUB (0/40,151,652 reads) and two *T. gondii* - COUGAR (0/31,562,801 reads) in BioProject: PRJNA61119 fails to identify any virus mapping reads. **C.** Ao contig 1 and contig 2 expression (reads per kilobase of million mapped reads, RPKM) and Pearson correlation between contig 1 and contig 2 expression across all mRNA and miRNAseq datasets in the Nĝo study, N = 237.

To quantify the relative abundance of human, parasite, and virus nucleic acids across each library, we aligned the reads against each respective genome and observed that high expression of the putative viral RdRp contig (RPKM *≥* 10) was specific to samples experimentally infected by the *T. gondii* - RUB strain (mRNAseq = 4/4, miRNAseq 5/5), and was mostly absent (RPKM *≤* 1.0) from other strains or controls (mRNAseq = 112*/*113, miRNAseq 115*/*115, Fisher’s Exact Test, p *<* 1*e−* 4) (Figure 2A). We rule out that the RdRp is an endogenous viral element since corresponding DNAseq data has zero virus-aligning reads. (Figure 2B). Trace RdRp expression (RPKM 0.5 - 2.0) was observed in three *T. gondii*-infected samples, all annotated as other strains: RH, GT1, and RH-GRA10[KO]. The relatively low abundance of sequencing reads is suggestive of sequencing cross-contamination (sequence batch information, supplementary Table S1), although early stages of viral cross-infection between samples cannot be ruled out, in which case the RUB strain would be a likely source (Figure 2C).

We propose that these two contigs constitute the genome of a bi-segmented RNA virus infecting *T. gondii* - RUB, which we name *Apocryptovirus odysseus* (Ao). The genus name derives from a Greek root suggesting a hidden or concealed virus, while we derive the species name from the leader of the soldiers who hid within the Trojan Horse in Greek myth, who when revealed wreaked havoc in the city of Troy.

### A. odysseus infects geographically distinct strains of T. gondii

Next, we investigated the global distribution of *T. gondii* and more broadly *Apicomplexa*-containing sequencing data and their association, if any, with Apocryptoviruses. We hypothesized that Ao would only be observed if its parasite host was present. To test this, we re-queried all 7.5 million sequencing runs processed by Serratus for high-similarity matches to Ao (*≥* 90% aa-identity) and identified one additional Ao+ BioProject (0.00052%, or 1/191,678 BioProjects), PRJNA114693. In the associated publication (“the Melo study”), the authors infected murine macrophages *in vitro* with 30 distinct strains of *T. gondii*, or a mock control (Melo et al., 2013). Following *de novo* assembly of all 32 Melo samples, we identified both Ao contigs in 2 samples: again a *T. gondii* - RUB strain (SRR446933), and *T. gondii* - COUGAR strain (SRR446909). Ao contig 1 and contig 2 exhibited a high expression (RPKM *≥* 10) in RUB and COUGAR libraries, and were absent (RPKM *≤* 1) from 28 other *T. gondii* strains or the mock control (Figure 3A).

**Figure 3.**
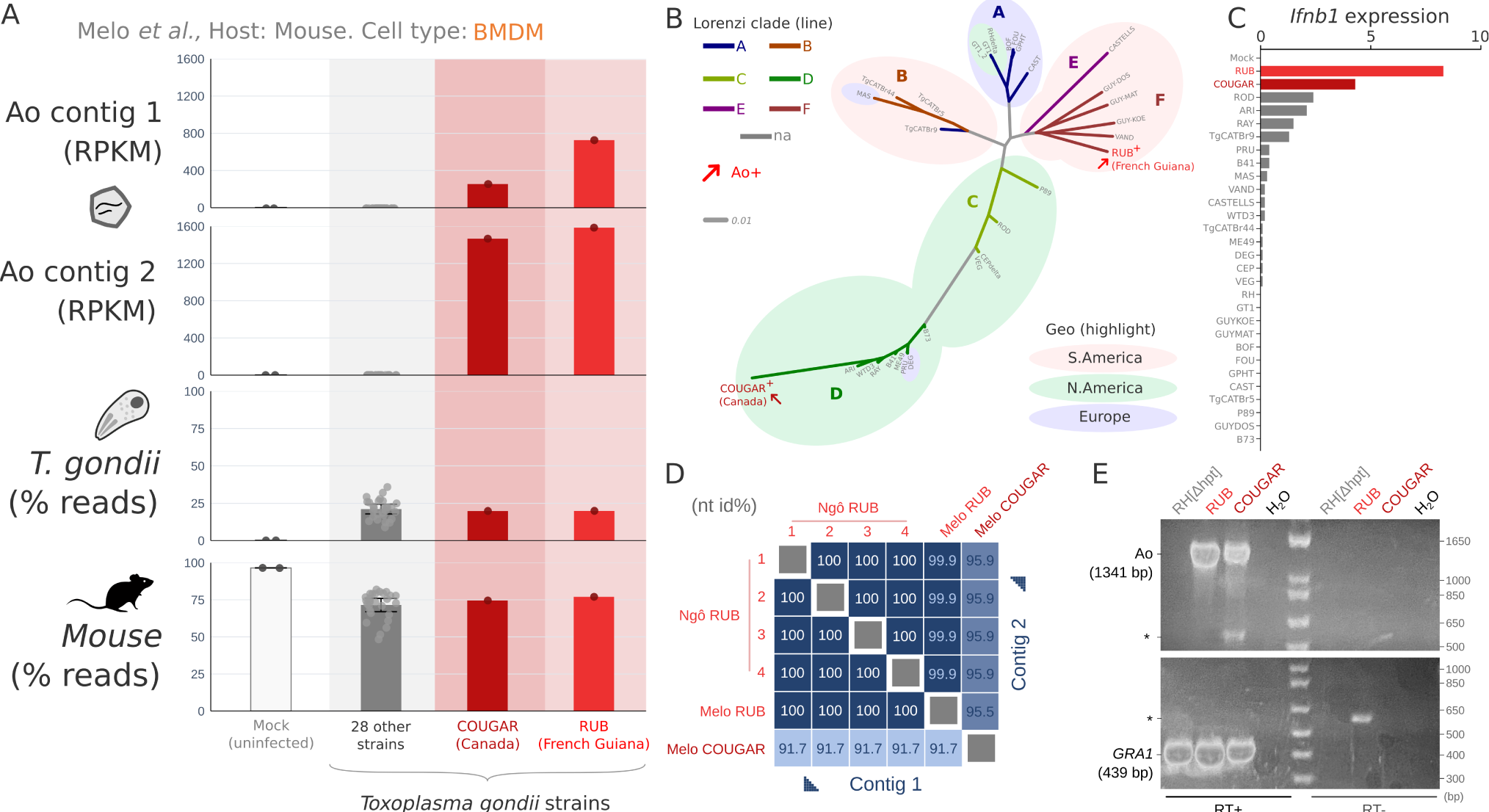
Quantification of *A. odysseus* in Melo *et al*. **A.** Quantification of Ao-aligning reads amongst 30 strains of *T. gondii* strains and mock control, error bars indicate 2 standard deviations. **B.** Unrooted *T. gondii* strain phylogeny (IQ-TREE) from transcriptome in the Melo study (149,777 measured sites; nodes have 100% bootstrap support, extended in Supplementary Figure S1), and strain cross-contamination ruled out by transcriptome variant analysis in Supplementary Figure S2. The original geographic isolation of strains is highlighted as North American (green), European (blue), or South American (pink), and lines are coloured by the genomic *T. gondii* clades defined by Lorenzi *et al*. (Supplementary Table S2). *A. odysseus* (Ao, red arrow) was detected at high levels in two geographically and phylogenetically unrelated strains of *T. gondii*; RUB isolated from a symptomatic soldier in French Guiana in 1991 and COUGAR isolated from a cougar in British Columbia in 1996. **C.** RNA sequencing expression data obtained from Supplementary Table 1 of the Melo *et al*. study showing normalized expression levels of *Ifnb1* in murine macrophages infected with strains of *T. gondii*. **D.** Pairwise nucleotide sequence identity of complete coding sequences from contig 1 and 2 from the Ngô and Melo studies. **E.** Reverse Transcriptase (RT) PCR for Ao or GRA1, a highly expressed Toxoplasma gene used as an RNA quality control, from Ao+ (RUB and COUGAR) and Ao-control RH[delta hpt]) strains of *Toxoplasma* grown on human foreskin fibroblasts. Asterix denotes non-specific bands. Sanger sequencing of amplicons showed 100% nucleotide identity to the assembled Ao RUB and Ao COUGAR, respectively.

To assess whether Ao positivity in RUB and COUGAR could be explained by genetic similarities between the two strains, we performed a phylogenetic analysis of the *Toxoplasma* parasite. Over the estimated 1 million years since *T. gondii* arose (Bertranpetit et al., 2017), the parasite has radiated into phylogeographically distinct clades A through F (Lorenzi et al., 2016). *T. gondii* - RUB and COUGAR strains are phylogenetically and geographically divergent from one another; RUB is a clade F strain isolated from a human in French Guiana (Dardé et al., 1998), while COUGAR is a clade D strain isolated from a mountain lion associated with a toxoplasmosis outbreak in British Columbia, Canada (Aramini et al., 1998) (Figure 3B), yet as noted by ”Melo” et al., these unrelated strains are both strong inducers of interferon beta (*Ifnb1*) (Figure 3C).

To identify additional RUB or COUGAR RNA sequencing data, we performed a metadata search but failed to find any additional datasets in the SRA from these strains. Given the available data, Ao appears to be fully penetrant amongst *T. gondii* RUB and COUGAR RNAseq libraries. Comparing the Ao RdRp coding sequences across samples, the Ao RUB from Nĝo and Ao RUB from Melo had 100% nucleotide identity (nt id), while the Ao RUB and Ao COUGAR had only 91.7% nt id. There is no read-level support for cross-contamination between *T.gondii* RUB and COUGAR samples (Figure S2), which combined with the cross-sample Ao virus sequence variation, supports that the Ao RUB and Ao COUGAR are distinct strains of the virus. Further, Ao viral RNA was confirmed by reverse transcriptase PCR (RT-PCR) in *T. gondii* cultured on human fibroblasts in the RUB and COUGAR strains, and were absent in RH[delta hpt] strain, and without the addition of RT (control) (Figure 3E).

Finally, we also performed a BLASTn search of the Ao COUGAR RdRp contig using the BLAST expressed sequence tag (est) database (accessed: 2024-02-22), and identified 15 ESTs (top hit: CV549349.1, mean nt id: 98.3%, max e-value: 2e-90, size range: 189 to 805 nt, sample submission date 2004-07-01. Query: Ao RUB, mean nucleotide identity: 90.4%) which were all isolated from *T. gondii* COUGAR tachyzoites (Ajioka et al., 1998). Likewise, searching for Ao COUGAR contig 2 in the est database revealed 72 ESTs (top hit: CV653441.1, mean nt id: 99.1%, max e-value: 8e-56, size range: 165 to 601 nt. Query Ao RUB, mean nt-id: 93.85%).

The available samples of RUB and COUGAR are experimental/laboratory-associated, therefore it is undetermined if Ao is found in wild isolates. Regardless, given the evolutionary, geographic source, and *T. gondii* host strain differences between the Ao RUB and COUGAR viral strains, and the absence of virus in intermediate *T. gondii* strains, it is plausible that Ao is a horizontally infectious virus circulating in *T. gondii*.

### Apocryptoviruses are a diverse clade of parasite-associated Narnaviruses

To elucidate the evolutionary and ecological context of Ao, we interrogated the viruses closely related to the virus. Although hidden Markov model (HMM) homology search via PFAM and InterProScan (Finn et al., 2014; Jones et al., 2014) failed to recognize Ao’s RdRp (date accessed: June 2023), remote homologs were identified with BLASTp within the *Narnaviridae* (date accessed: June 2023). This discrepancy is likely due to a lack of adequate narnavirus representation in standard HMMs: Ao shares only 25% and 28% amino acid identity with the exemplar Narnaviruses ScNV-20S and ScNV-23S, respectively. The highest similarity match to Ao RdRp was an unclassified ribovirus partial RdRp (52% aa identity, query coverage 40%, subject coverage 100% accession: QIM73983.1) isolated in 2016 from the lung of a Ryukyu mouse (*Mus caroli*) in Thailand (Wu et al., 2021). The remaining Narnavirus matches were distal, at below 40% amino acid identity.

Due to a paucity of data within the family *Narnaviridae*, and the divergent nature of Ao, we re-queried Serratus for Ao-like RdRp palmprints. We retrieved and assembled 166 matching runs from 46 BioProjects with 32 species-like operational taxonomic units (sOTUs) (186 distinct virus-run observations, Supplementary Table S3) (Chen et al., 2022; Zhou et al., 2021; Gil and Hird, 2022; Gao et al., 2021; He et al., 2021; Chen et al., 2021b; Rhie et al., 2021; Bittner et al., 2022; Chen et al., 2021a; Xie et al., 2021; Cui et al., 2022; Wen et al., 2022; Zhou et al., 2022b; Sun et al., 2022; Wu et al., 2021). We recovered relevant Narnavirus RdRp contigs for all 32/32 (100%) sOTU, of which 22/32 (75%) contained a sufficiently long RdRp sequence (*≈* 390 amino acids, motif F-E (Venkataraman et al., 2018) and thumb) for robust phylogenetic reconstruction. In addition, we enforced that RdRps must be *<* 90% identity from other members to be designated as distinct species, otherwise sequences were considered as strain-level variants. Thus, two pigeon-associated sOTUs (*Columba livia*) were considered as one virus species *A. anticulus*; *A. pancratius* described here is a strain of the *Ribovirus sp.* (RtMc-NcV/Tu2016) (QIM73973.1) and four domestic animal-associated sOTUs (two *Ovis aries*, *Capra hircus*, and *Sus scrofa*) were considered as one species: *A. demophon*. In total this yields 20 species members of the genus *Apocryptovirus* Figure 4 and Extended Table S3.

**Figure 4.**
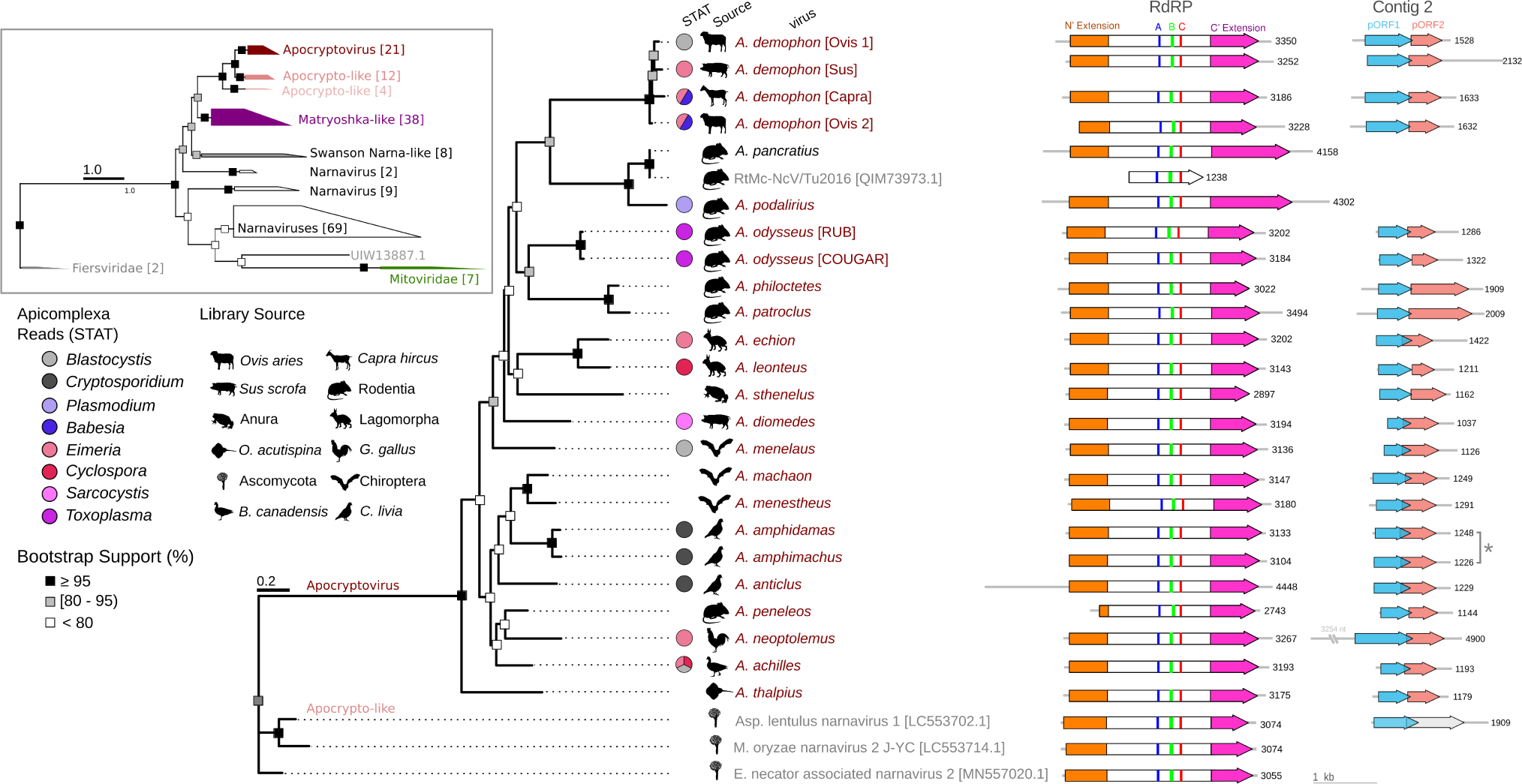
Apocryptovirus RdRp phylogeny and genome maps. Maximum-likelihood phylogenetic tree (IQ-TREE, 2.1.4& ggtree, 3.10.0) and contigs 1 and 2 for the 20 proposed species of *Apocryptoviruses* estimated based on RdRp palm (motifs F-E) and thumb subdomains, with an outgroup of representative Apocrypto-like viruses. **Inlay**, shows the placement of *Apocryptoviruses* within *Narnaviridae* (see: Figure S4). Novel viruses are written in red and GenBank viruses in gray. Scale bars represents amino acid substitutions per site, and square symbols denote bootstrap confidence. For each *Apocryptovirus*, the source sequencing library taxonomic label (silhouettes), and co-occuring *Apicomplexa*-categorized reads (STAT (Katz et al., 2021)). Conserved RdRp ORF (contig 1) and pORF1,2 contigs with contig length indicated in nucleotides. *A. amphidamas* and *A. amphimachus* were recovered from a common sequencing library, therefore the corresponding contig 2 is not known with certainty (denoted with asterisk).

Using both the novel RdRps recovered from the SRA and alignable RdRp hits in GenBank, we constructed a maximum-likelihood phylogenetic tree (Figure 4). Ao and the Ao-like Serratus hits, as well as the rodent lung virome Narnavirus, form a monophyletic clade (bootstrap support 100%). These RdRp share at least 70% amino acid identity with another member in the clade, and less than 70% identity with all members outside the clade (Supplementary Figure S3). We propose that these viruses, together with Ao, constitute a novel genus within *Narnaviridae*. Apocryptoviruses are situated within a larger Apocryptovirus-proximal clade that also comprises the Matryoshka RNA viruses (Charon et al., 2019), all of which are in *Narnaviridae* with high confidence (Figure 4). Individual viral species of this new genus have been named after the other soldiers hidden inside the Trojan Horse that were led by Odysseus in the myth.

The prevalence of Ao and the Apocryptoviruses is rare amongst SRA samples. Only 6/23,530 (0.025%) runs annotated as *T. gondii* are Ao-positive, and zero runs containing *≥* 1024 *Toxoplasma sp.* reads measured by STAT (Katz et al., 2021), and not annotated as *T. gondii*, are Ao-positive (0/9,071). The rate of Ao-positivity is estimated to be below 1% in *Toxoplasma*.

Predicting virus host range based on metagenomic data is an ongoing challenge in viromics. The overwhelming majority of *Narnaviridae* are metagenomic or from complex samples such as leaf lesions. The closest identified Narnavirus with a plausible host assignment is Aspergillus lentulus Narnavirus 1 (AleNV1, BCH36643.1), isolated from cultured *Aspergillus lentulus* mycelia (Chiba et al., 2021). The next closest are the Matryoshka RNA viruses isolated from human blood co-infected with *Plasmodium* or *Leucocytozoon* parasites (Charon et al., 2019).

Nominally, Ao was identified in libraries labelled as *H. sapiens* (Nĝo et al., 2017) or *M. musculus* (Melo et al., 2013); yet text metadata and taxonomic classification of nucleic acids in those libraries pointed us towards *T. gondii* as the common virus-associated factor, suggesting the parasite was the host for Ao. This virus-host relationship was strengthened when Ao was found in multiple replicates specific to 1/9 *Toxoplasma* strains in the Nĝo study, and 2/31 strains in the Melo study, as well as in *Toxoplasma* ESTs. Generalizing this rationale, we decided to measure the extent of apicomplexan positively in each of the Apocryptovirus-positive libraries and found that 10/18 (55%) of the viruses were associated with *Apicomplexa* in at least one dataset (Figure 4 and Supplementary Table S3).

To further test the relationship between *Apocryptovirus* and *Apicomplexa* we calculated their rate of co-occurrence per library for each source organism and compared this to the background rate of *Apicomplexa*-positively. We found that apocryptoviruses are highly predictive (Fisher’s exact test) for the co-occurrence of *Apicomplexa* in pig (p = 2.42E-09), chicken (p = 2.34E-14), sheep (p = 2.02E-18), goat (p = 3.64E-13), rabbit (p = 3.09E-15), and sparrow (p = 5.27E-05) (Figure 5). Conversely, we measured the prevalence of apocryptoviruses amongst Apicomplexa-positive samples in the SRA and found the viruses occurred variably in pig (6/447, 1.34% virus-positive), chicken (8/445, 1.79%), sheep (24/444, 5.40%), goat (9/151, 5.96%), rabbit (11/61, 18.03%), and sparrow (5/12, 41.66%) samples.

**Figure 5.**
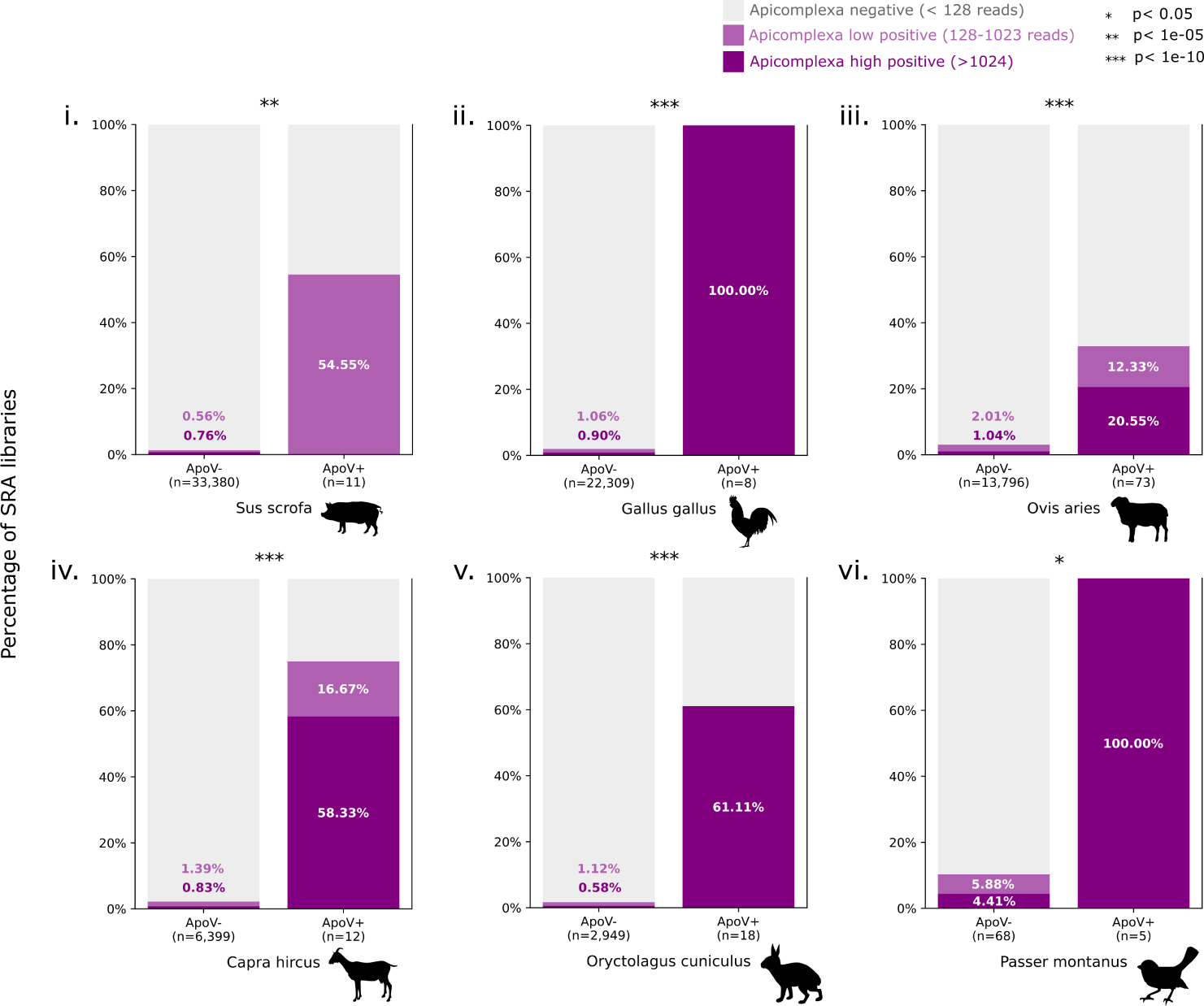
Apocryptoviruses are predictive of Apicomplexa co-occurence. Samples from six non-laboratory animals (i. pig, ii. chicken, iii. sheep, iv. goat, v. rabbit and vi. tree sparrow) were first categorized as Apocryptovirus-negative (ApoV-) or Apocryptovirus-positive (Apo+). We then measured the amount of STAT classified Apicomplexa-reads in each library and classified each library as; Apicomplexa-negative, *≤* 128 reads (gray); low Apicomplexa-positive, 128-1023 reads (light purple); or high Apicomplexa-positive, *≥* 1024. The significance of the co-occurrence was evaluated with a Fisher’s Exact Test (Methods), for *Apocryptovirus* +/- and *Apicomplexa*(high or low)+/-.

Parsimony suggests apocryptoviruses are infecting *Apicomplexa*, which in turn would allow the viruses to enter, but not infect, vertebrate host cells. Currently there is no evidence that apocryptoviruses can replicate in vertebrates in the absence of an apicomplexan. As such, in any molecular interaction between a virus-infected apicomplexan and the macrohost, the virus is likely a bystander. An alternative hypothesis is that apicomplexan infection sensitizes the vertebrate cells to apocryptovirus infection/replication, but the resolution of these hypotheses will require further experimental validation.

The viruses *A. neoptolemus*, *A. diomedes*, and (particularly) *A. demophon* showed a high prevalence in parasites associated with livestock, and may thus modulate the biology of agriculturally-relevant pathogens. Of note among these is *Eimeria sp.*, a causative agent of coccidiosis, a high mortality disease causing billions of dollars of economic loss for farmers (Blake and Tomley, 2014).

### Structural characterization of Apocryptovirus proteins

The structure of narna-like RdRp has not been well-characterised. Comparing the predicted structure for Ao RUB RdRp (confident, predicted local distance difference test: 84.56) to the experimentally-solved Poliovirus RdRp structure, Ao RdRp folds into a palm superfamily structure with fingers, palm and thumb (Figure 6) with intact RdRp sequence motifs (Venkataraman et al., 2018). We confirmed that the *Apocryptovirus* contig 1 encodes a biochemically competent RdRp on the basis of core RdRp motif conservation (Figure 6A). We noted that the RdRp of Apocryptoviruses have N’- and C’-extensions of 220 and 224 amino acids long, respectively, that extend beyond the palm, fingers, and thumb of a minimally-viable RdRp (with Poliovirus being a well-studied exemplar of a minimal RdRp). To investigate the possible activity of these extensions, we first sought to analyze sequence motifs conserved in the Apocrypto-proximal clade, then apply these to structure. We re-aligned more closely-related (Apocrypto-proximal) RdRps, which allowed us to create a higher confidence multiple sequence alignment (MSA) of the full protein sequence, inclusive of the extensions. We identified 12 high-occupancy and highly conserved regions from the MSA and designate them with lowercase Greek letters *α* through *µ* (Figure S5).

**Figure 6.**
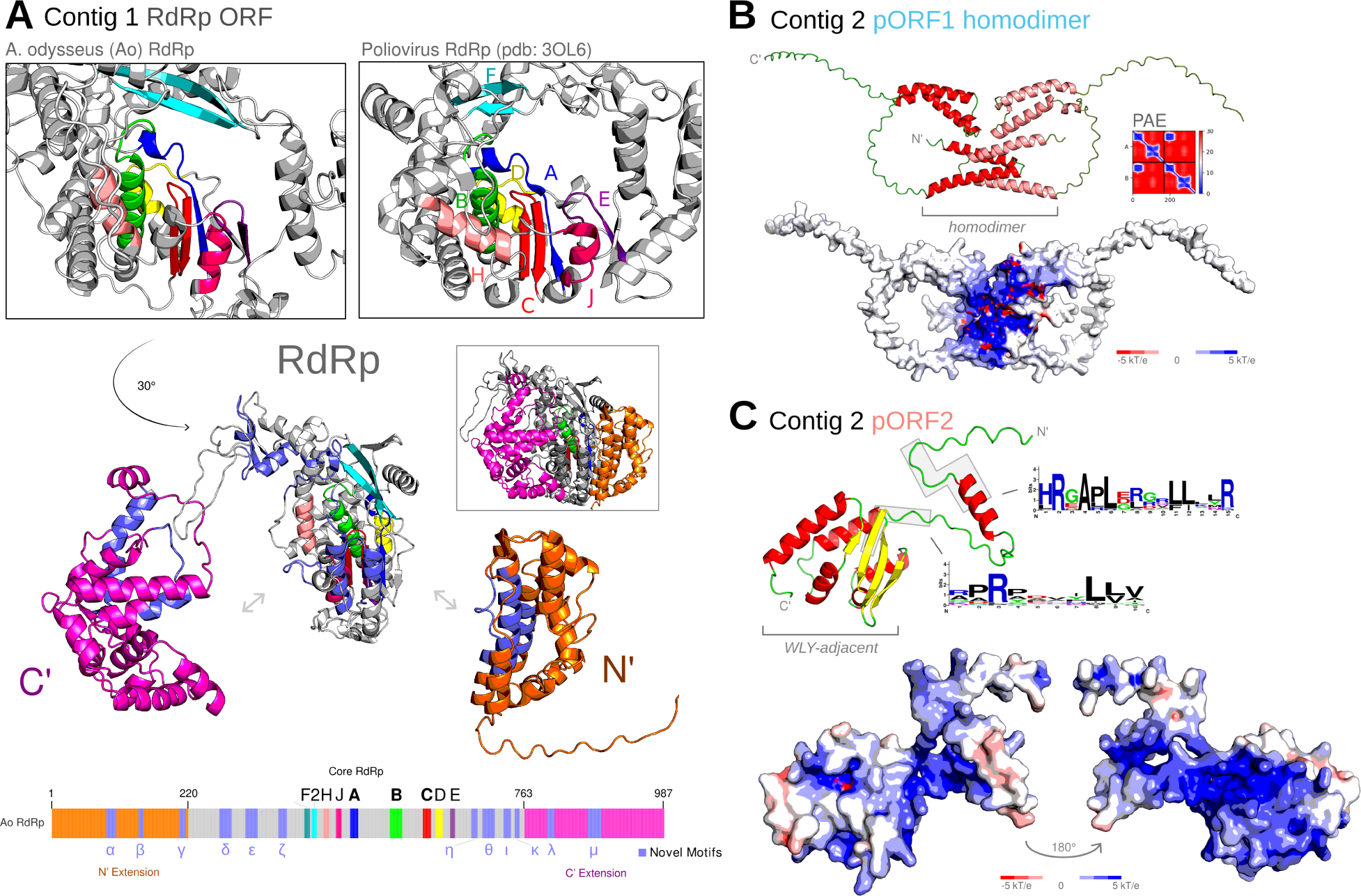
Protein Structural Analysis. **A.** Predicted structure of Ao RUB RdRp and solved X-ray structure of poliovirus RdRp (PDB accession 3OL6 (Gong and Peersen, 2010)), at their respective catalytic cores, with motifs F-E highlighted (Venkataraman et al., 2018). Ao RdRp is shown in an exploded view, rotated with motif C facing forward. The N’ and C’ terminal extensions form distinct domains from the catalytic core RdRp. (Extended Figure S6). The Apocryptovirus sequence motifs had similar conservation to the RdRp core motifs against a background of sequences falling outside of the motifs — high occupancy inter-motif sequences (Extended Figure S5). **B.** Predicted structure of Ao RUB contig2 pORF1 as a homodimer, and calculated surface electrostatic potential map (apbs v3.4.1 (Jurrus et al., 2018) in PyMOL v2.5.0 Open Source (Schrodinger, LLC, 2015)). The predicted aligned error (PAE) shows a high confidence interaction between the N’ alpha helices. **C.** Predicted structure and electrostatic potential map of Ao RUB contig2 pORF2, with conserved region sequence logo highlighted.

We reconstructed the MSA for the Ao RUB pORF1 and pORF2 (HMMER, 3.3.2) to improve protein structure prediction with ColabFold. The pORF1 has confident predictions for two positively-charged helix-turn-helix pairs with intervening disordered loops. We then assessed if this protein could form homodimers, and indeed the positive surface charges localize to one another, and the pair of helices interlock in a putative homodimerization domain Figure 6. The pORF2 was predicted to be more structured, with an N’ alpha helix coinciding with a conserved positive-charge motif, and a short unknown domain. The linker and alpha-helices of pORF2 have a positively-charged surface with well-conserved Arg residues Figure 6. Searching for structures similar to pORF2 with FoldSeek (accessed: 2024-02-24) (van Kempen et al., 2023), uncovered 15 hits (probability *≥* 0.95, e-value *≤* 1e-1), 13 (86.7%) of which are annotated as being WYL domain-containing. This Ao pORF2 domain is itself not a WYL-domain, but pORF2-like domains are found adjacent to WYL. WYL domains can bind nucleic acids and are well characterized to play a role in transcriptional regulation in bacteria (Keller and Weber-Ban, 2023); considered together with the conserved positive charges on pORF2, we hypothesize this protein is involved in a nucleic acid interaction.

Related Narnaviruses have reported second segments: Aspergillus lentulus Narnavirus 1 (Chiba et al., 2021), Matryoshka viruses (Charon et al., 2019), and Leptomonas seymouri Narna-like virus 1 (a PPV) (Sukla et al., 2017), although BLASTp did not retrieve this as a match. To test for remote homology we constructed HMMs for *Apocryptovirus* pORF1 and pORF2. The pORF1, but not the faster evolving pORF2, (Figure S3B), models matched the Aspergillus lentulus Narnavirus 1 ORF1 (hmmscan v3.4, E-value: 4.6e-10, bitscore 26.6) establishing a remote homology between the contig 2 of these related Narnaviruses.

### Is A. odysseus a T. gondii hypervirulence factor?

*T. gondii* RUB and COUGAR are atypically hypervirulent. RUB was isolated from an immunocompetent soldier in French Guiana (1991) presenting with fever, myalgia, and leukopenia which developed into rales, respiratory failure, and renal deterioration (Dardé et al., 1998). While the COUGAR strain was isolated from a mountain lion (Dubey et al., 2008; Dubey, 2021b), it is believed to be identical to the strain which infected 3000-7000 people in a water-borne 1994/1995 toxoplasmosis outbreak in Victoria, BC (Bowie et al., 1997). Notably, the incidence of ocular inflammation (retinitis) toxoplasmosis amongst these immunocompetent patients was high in the Victoria outbreak (Bowie et al., 1997). The Melo *et al*. publication notes that exactly these two “atypical” *T. gondii* strains are outliers by their capacity to induce a type I interferon inflammatory response in murine cells (see also: Figure 3C). We sought to re-analyse the Nĝo *et al*. human dataset, hypothesizing that the presence of Ao in both RUB and COUGAR provides a plausible mechanism for an immune-mediated hypervirulence of *T. gondii*.

Using the Nĝo *et al*. datasets, we performed differential gene expression analysis (DGE) and gene set enrichment analysis (GSEA) to test if the human immune response to Ao+ *T. gondii* strains was similar to the murine macrophage response from Melo *et al*., characterized by upregulation of the type I interferon gene *Ifnb1*. The Nĝo *et al. T.gondii* - RUB infection RNAseq data was available in two cell types, macrophage (n = 2) and neuronal stem cells (NSCs, n =2). We quantified human *IFNB1* induction in neuronal (Figure 7) and macrophage cell types (Supplementary Figure S7) across all *T. gondii* or mock datasets. We noted a large variation in *IFNB1* gene expression, especially across the macrophage datasets including a second set of mock controls, which could be explained by a sequencing batch-effect in the data (Supplementary Figure S7 and Supplementary Table S1). Samples segregated by batch in a principal component analysis (Supplementary Figure S8A) are indicative of global profile differences, thus we limit differential expression analysis to intra-batch comparisons.

**Figure 7.**
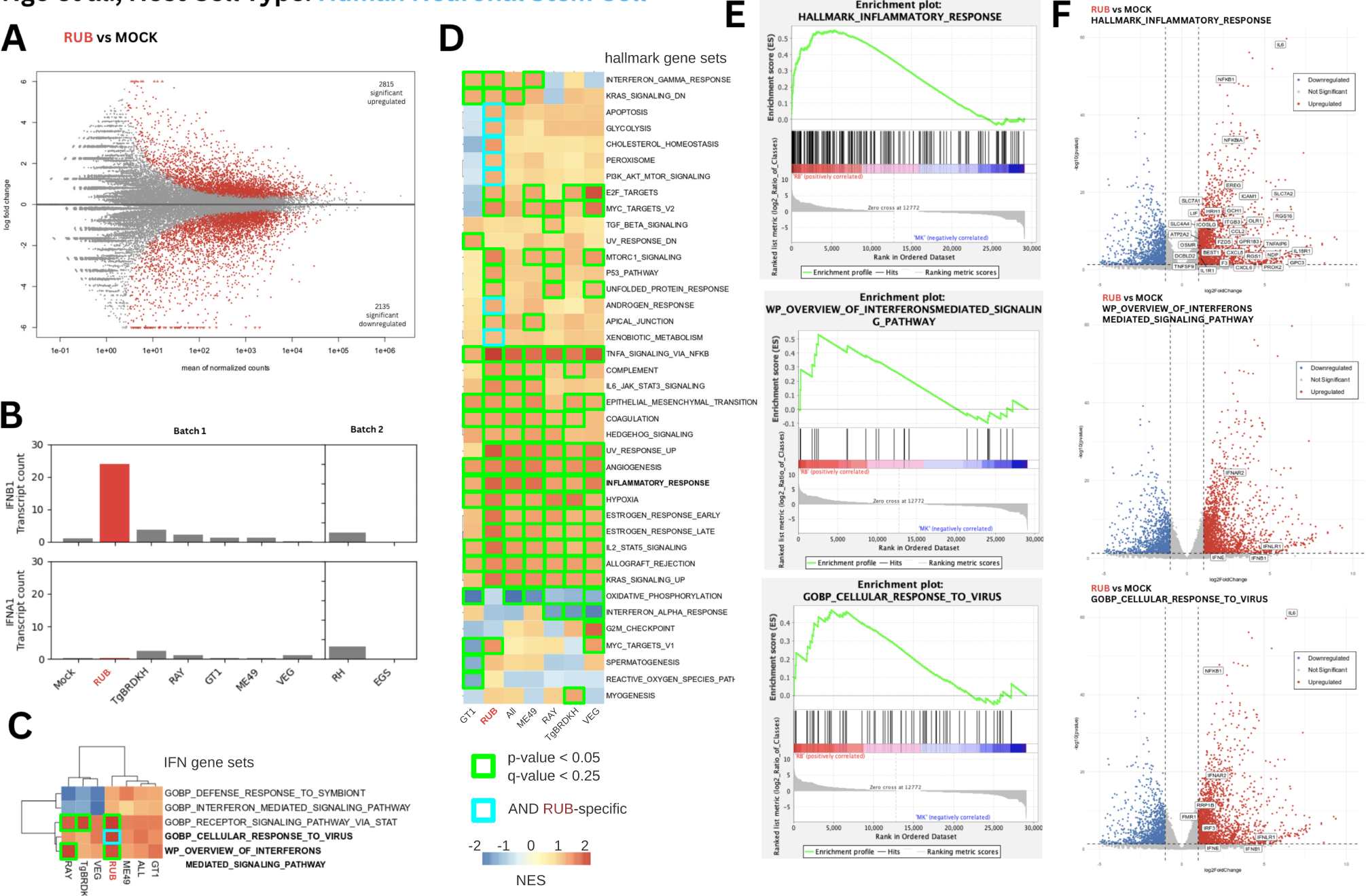
Differential Gene Expression (DGE) of human neuronal stem cells infected with various *T. gondii* strains. **A.** MA plot of *T. gondii* - RUB vs. Mock genes (highlighted: Benjamini-Hochberg adjusted p-value *<* 0.05). **B.** Bar plot of normalized transcription counts of *IFNB1* and *IFNA1* across *T. gondii* strains and Mock sequence in Nĝo *et al*. experiments, separated by batch. **C.** Heat map of Normalized Enrichment Scores (NES) from Gene Set Enrichment Analysis (GSEA) using gene sets possessing interferon-specific genes, namely *IFNA1* and *IFNB1*, applied to the *T. gondii* strains. **D.** Heat map of NES values from GSEA using the Hallmark gene sets, only significant gene sets are shown. **E.** GSEA curves comparing RUB vs. Mock strain using gene sets of inflammatory response, cellular response to virus, and interferon-mediated signalling pathway. **F.** Volcano plot of differentially regulated genes with significant members of notable gene sets being labelled.

Amongst the comparable *T. gondii*-infected neuronal samples, the RUB strain (n = 2) induced the highest *IFNB1* expression (26.1 fold increase vs. Mock; n = 2, p = 0.029) relative to TgBRDKH (n = 2), GT1 (n = 2), ME49 (n = 2), RAY (n = 2) or VEG (n = 2) (Figure 7B), in agreement with the murine *Ifnb1* expression in the Melo study (see also Figure 3C). Next, we sought to contextualize the type of *IFNB1* response, asking which of the five gene sets containing *IFNB1*, if any, are differentially regulated in RUB versus mock and the rank of this difference amongst other *T. gondii* strains. While infection with both RUB and RAY strains show upregulation of the gene signature for “Interferon mediated signalling pathway” (p-value 0.009 and 0.049) and “Receptor signalling via STAT”, (p-value 0.006 and 0.039) a known downstream effector of interferon, the RUB strain was specifically enriched for “cellular response to virus” (p-value 0.019), supporting the existence of specific host biological viral-response against Ao.

To characterize the broader host response of *T. gondii* RUB, we performed GSEA using the well-defined hallmark gene set (Liberzon et al., 2015). As expected, all *T. gondii* infections induce an “Inflammatory Response” with the involvement of the “TNFA signalling via NF*κ*B” pathway relative to mock controls. Yet the magnitude of these responses in neurons is the highest in RUB when compared to other (non-viral) strains (Figure 7D). In addition, RUB was the only *T. gondii* strain inducing “Apoptosis”, “Glycolysis”, “Peroxisome”, and the “PI3K AKT MTOR Signalling” axis, which altogether is consistent with the Melo study conclusions that *T. gondii* RUB is exceptionally pro-inflammatory, even amongst strains of *T. gondii*.

A similar inflammatory response trend is observed in the *T. gondii* RUB macrophage-infection experiments, but the response is less specific/exceptional (Supplementary Figure S7). A higher overall level of experimental variation in the macrophages (n = 2 each) is evident when comparing the MA plots across cell types for statistically differentially regulated genes (compare NSC Figure 7A and macrophages Supplementary Figure S7A).

A major limitation of these *in silico* analyses is that we cannot establish a causal relationship between Ao and the host inflammatory responses. Notably, *T. gondii* RUB genetic factors are likely to confound this analysis. Yet we are able to recapitulate a statistically significant association of the Ao+ RUB strain with an inflammatory response in human cells. In the Melo *et al*. study, this inflammatory response was experimentally demonstrated to be nucleic-acid mediated which, taken together with the discovery of Ao, allows us to tentatively propose a virus-mediated hypervirulence model (Figure 8).

**Figure 8.**
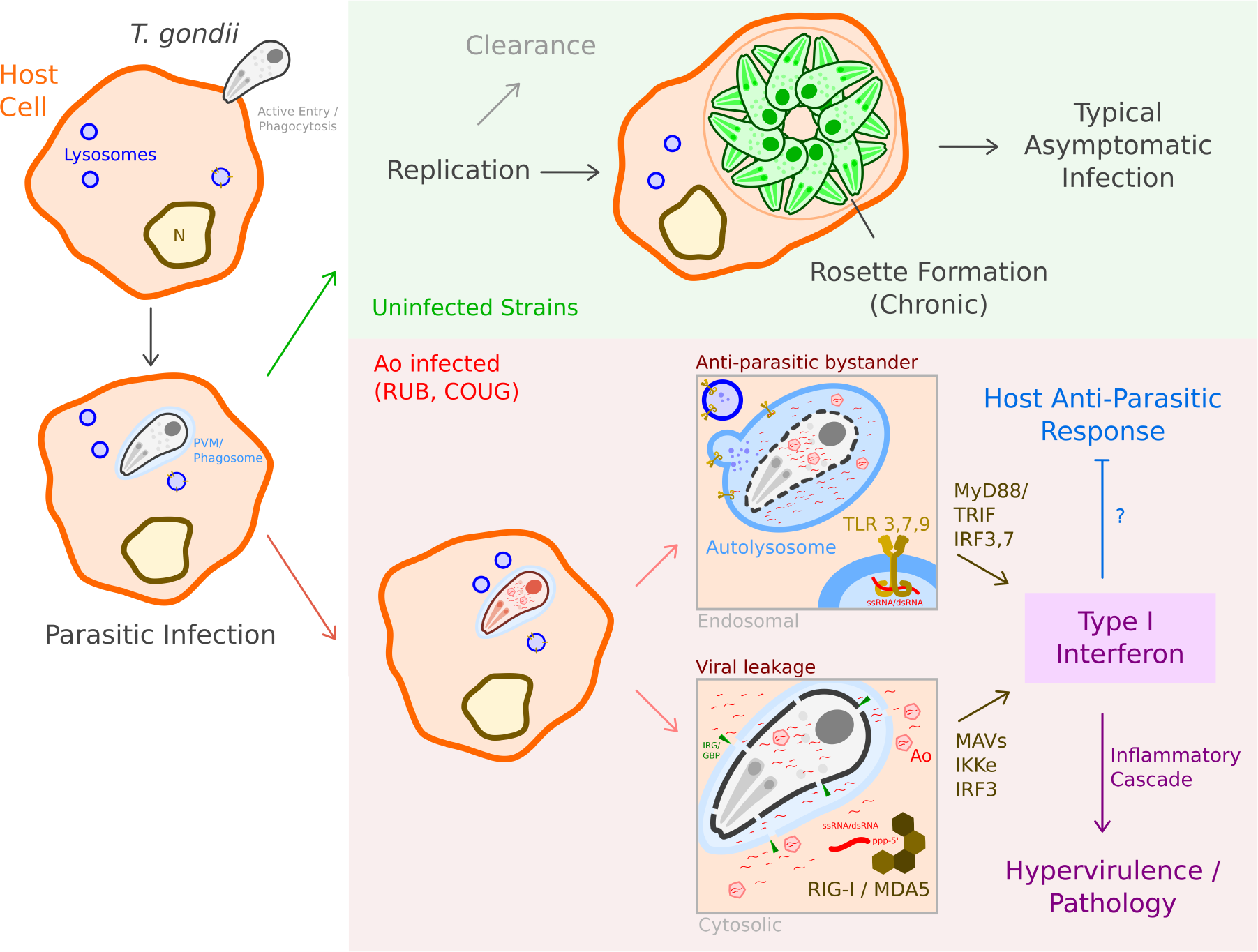
Hypervirulence model *T.* *gondii* infects host cells. In uninfected strains, this either leads to successful clearance by the host or the establishment of chronic infection, typically asymptomatic. In virus+ strains, such as RUB and COUGAR, detection of viral RNA (vRNA) - either by endosomal TLRs after phagolysosome maturation following phagocytosis, or by RLRs surveying the cytosol after interferon-inducible GTPase (IRG/GBP)-mediated destruction of the parasitophorous vacuolar membrane - leads to the induction of type I IFN. The effect of this is twofold - the viral pathogen-associated molecular pattern drives additional inflammation which leads to pathology and overt symptoms, but potentially also impairs or diverts host anti-parasitic immunity. IRG: Immunity Related GTPase; GBP: Guanylate Binding Protein.

## Discussion

We discover a novel narnavirus, *Apocryptovirus odysseus*, which tightly associates with two distinct yet hypervirulent strains of *T. gondii* - RUB and COUGAR. Ao is a member of a broadly distributed clade (putatively at the genus level) of narnaviruses, the *Apocryptovirus*. We also describe 18 additional novel species in *Apocryptovirus*, which likely infect apicomplexans which in turn infect chordates. Furthermore, we provide initial (*in silico*) evidence to assert that Ao within *T. gondii*, may act upon the innate immune system of human and mouse cells and therefore may be a hitherto uncharacterized hypervirulence factor for this ubiquitous parasite.

While PPVs have been described for decades, their entanglement in the parasite-host relationship and pathogenesis is more recent (Gömez-Arreaza et al., 2017; Zhao et al., 2023). Multiple causal studies establish a molecular mechanism by which parasite-infecting viruses trigger chordate innate immune responses. Examples of this menage à trois include TVV in *Trichomonas vaginalis* (Fichorova et al., 2012), Leishmania RNA virus 1 (LRV1) in *Leishmania Viannia sp.* (Ives et al., 2011; Eren et al., 2016), GLV1 in *Giardia lamblia* (Pu et al., 2021) and CSpV1 in *Cryptosporidia sp.* (Deng et al., 2023). Unlike Ao, previously-characterised PPVs are double-stranded RNA viruses, and a narnavirus contributing to disease in vertebrates has not yet been characterized. A recurrent theme amongst these studies is enhanced host pro-inflammatory cytokine production, specifically interferon, which can worsens parasite virulence.

Given the recurrent examples of PPV-mediated pathogenesis and the scarcity of examples of narnavirus infection-related phenotypes in the viral host, it is parsimonious to suggest that the presence of Ao RNA is a sufficient trigger of macrohost IFN-driven immune responses (Figure 8). Melo *et al*. found that induction of type I IFN in the COUGAR strain is dependent upon the RNA-sensor RIG-I, and the signal cascade proceeds through MyD88/TRIF and ultimately IRF3, an inducer of type I IFN (Melo et al., 2013). IRF3 phosphorylation and IFN-induction are also observed in other PPV infections, albeit with a double-stranded RNA ligand (Ives et al., 2011; Fichorova et al., 2012; Eren et al., 2016; Deng et al., 2023). Not knowing of the existence of Ao within their experiment, the authors ascribe COUGAR/RUB hypervirulence to parasite genomic DNA/RNA, potentially associated with the enhanced killing observed in human foreskin fibroblasts, leading to release of greater quantities of *Toxoplasma* nucleic acids (Melo et al., 2013). This is plausible and not necessarily inconsistent with a contribution of vRNA to the innate response, although whether similar differences in killing can also be seen after phagocytosis by antigen-presenting cells such as macrophages (a fundamentally different process to infection of a fibroblast, during which the *Toxoplasma* mediates its own entry into a parasitophorous vacuole) is unclear. It also cannot explain the specificity of type I IFN response in these two strains and absence from the remaining 29 (Ao- and ApoV-) strains which also possess parasite nucleic acids (Melo et al., 2013). With this in mind and in the absence of similar enhanced killing data for the RUB strain, we find that the presence of Ao in both strains remains the single most persuasive explanation for the differential host immune response. We also note that Melo *et al*. find significant heterogeneity between the transcriptomic profile of both murine macrophages responding to infection by these two strains and indeed between the two strains themselves. The only obvious transcriptomic similarity is the induced type I IFN response, which can be neatly explained by the key common denominator which separates the two otherwise unrelated COUGAR and RUB strains from all others - Ao positivity.

We do note differences between existing pathways previously elucidated for innate response to PPVs and a potential contribution of Ao to pathogenicity. The other cases involve dsRNA viruses via TLR3/IRF3, whereas a ssRNA narnavirus genome would likely trigger TLR7, 9 (ssRNA ligand) or a RIG-like receptor (uncapped RNA), and downstream of this, display some dependence on *IRF7* (and, had it been investigated, *IRF5*) (Jensen and Thomsen, 2012; Schoenemeyer et al., 2005). Melo *et al*. however found that murine macrophages infected by COUGAR strain tachyzoites, *Irf7[KO]* in fact increases *Ifnb1* expression (Melo et al., 2013). This discrepancy could arise due to pleiotropic effects in constitutive immune-gene knockouts, and will additional experimental evidence to be resolved.

There is good evidence to suggest that the type I IFN response plays an important role in the control of *Toxoplasma*, even if debate continues around the direction of this response (i.e. whether type I IFN limits or supports infection). Knockout of *IFNAR* appears protective and supplementation of type I IFN detrimental after intraperitoneal injection of PRU strain *Toxoplasma* in mice (Hu et al., 2022). Similarly, an intestinal epithelial cell-specific knockout of *Ifnar* is protective in mice after oral infection of *Cryptosporidium* (another apicomplexan pathogen) carrying CSpV1 (Deng et al., 2023). In *Leishmania donovani* infection, *Ifnar* ^-/-^ mice have significantly lower parasite burden compared to wild-type (Kumar et al., 2020). The authors propose that type I IFNs inhibit dendritic cell activity that would otherwise prime a type II IFN-driven, CD4^+^ Th1 cell-mediated anti-parasitic defence; they also demonstrate that ruxolitinib, a small molecule inhibitor of JAK kinases involved in interferon transduction, exerts anti-parasitic effects that are not observed in *Ifnar^-/-^*mice, suggesting a dominant effect via inhibition of type I IFN signalling (Kumar et al., 2020).

On the other hand, others have found that after oral infection of ME49 cysts, *Ifnar^-/-^* mice had significantly poorer survival rates and increased numbers of brain cysts (Matta et al., 2019). Differences between strains, route of infection, and multiplicity of infection, with effects on the timing, duration, and magnitude of the type I IFN signal, could explain these contrasting findings (Silva-Barrios and Stager, 2017; Lee and Ashkar, 2018). Regardless of the eventual outcomes on parasite burden and host survival, we propose that the additional type I IFN response, brought on by the presence of Ao vRNA, will at least acutely contribute to more severe disease in COUGAR and RUB strain infection. While it has been suggested that PPVs may promote pathology driven by their host via immune modulation of the macrohost, such as has been recently observed in *Cryptosporidium* (Deng et al., 2023), we do not expect this to hold true across all parasite strains, species and hosts; the virus-parasite-host fitness relationships are by definition complex.

If it is indeed the case that PPVs cause exacerbated immune response in parasitic infection, it may be worthwhile to investigate antivirals as a form of treatment. This has already been suggested in the case of TVV+ *T. vaginalis*, wherein the canonical anti-parasitic, metronidazole, can counterintuitively exacerbate inflammatory signalling in infected epithelia (Fichorova et al., 2012; Narayanasamy et al., 2022). Elimination of LRV1 by RdRp inhibition resulted in the generation of strains which induce weaker inflammatory responses and were associated with reduced pathology *in vivo* (Kuhlmann et al., 2017). Similar investigations have focused on the efficacy of antivirals indinavir and ribavirin, individually or in combination with the anti-parasitic paromomycin against CSpV1+ *Cryptosporidium* in a 2D monolayer model of human enteric epithelia (Deng et al., 2023). Efforts during the COVID-19 pandemic highlighted the RdRp as an important therapeutic target in RNA virus antiviral therapy (Tian et al., 2021). Given the potential dependence of the efficacy of such drugs on RdRp structure, further elucidation of the structure in the *Narnaviridae* and proximal clades, using conventional wet lab approaches and molecular docking software, may build on our observations of conserved residues in Ao and related viruses to support the identification of effective antivirals. Besides, the generation of virus-free parasite strains will allow for elucidation of the contribution of endogenous virus to host and macrohost physiology (Kuhlmann et al., 2017; Espino-Vázquez et al., 2020; Narayanasamy et al., 2022; Kuroki et al., 2023).

While *T. gondii* pathogenicity is certainly multi-variate, with host immune status, age, and parasite genetic variation known influences on pathophysiology, there is still a lack of reliable prognostic indicators. A viral hypervirulence factor can explain, at least in part, some of the stochastic nature of parasitic disease burden in humans and other animals. Given the enormous infectious burden of *T. gondii* and other parasites (impacting multiple billions of people), it is prudent to improve monitoring for viruses within the niche of the human biome, as it interlinks human health with the health of microbial flora. In general, if the presence of PPV is prognostic (as it is in TTV/CSpV1), then ApoV+/- status can impact treatment and outbreak-management decisions in toxoplasmosis and other parasitic diseases.

Hypervirulence in *Toxoplasma* can present with different symptoms. Broadly, however, we note that in humans, both infections with COUGAR or RUB are associated with ocular involvement and overt acute pathology even in immunocompetent patients (Bowie et al., 1997; Dubey, 2021b; Dardé et al., 1998). In mice, both strains are associated with increased mortality (Khan et al., 2009; Niedelman et al., 2012; Hassan et al., 2019). COUGAR is associated with enhanced oral infectivity when compared to other strains, although whether RUB shares this phenotype is unknown (Su et al., 2003). A recent report of COUGAR strain infection in four southern sea otters (*Enhydra lutris nereis*) observed severe myocarditis and marked subcutaneous and peritoneal steatitis (Miller et al., 2023). Type I IFN signalling is thought to play an important role in driving adipocyte inflammation (Chan et al., 2020), so steatitis could present an unusual consequence of the vRNA contribution to toxoplasmosis in Ao+ strains.

Ultimately, these proposed mechanisms and models require experimental validation, for which we outline an efficient set of critical experiments. First, necessity and sufficiency of Ao-mediated IFN production/pathogenicity needs to be established. For necessity, Ao infection of a naive *T. gondii* strain (ME49) will promote *IFNB1* expression relative to an uninfected control. Curing of COUGAR/RUB *T. gondii* of the virus (for example, by serial passage in antiviral-supplemented culture until Ao-status can be confirmed by PCR) will cause a loss of IFN production/pathogenicity (sufficiency), and this effect can be rescued by subsequent re-infection of the cured strain. The gain and loss of type I IFN responses will be dependent on ssRNA receptors TLR7,9 or RLRs, and not TLR3 or TLR11,12. These critical experiments when performed *in vitro* would establish a causal relationship between Ao and proinflammatory induction, while the same set of experiments in an animal model can formalize the relationship between Ao and parasite pathogenicity.

The objective of this study was to rapidly screen for a candidate highly divergent neuroinflammatory virus *in silico*. Here we discovered *Apocryptovirus odysseus*, a narnavirus likely infecting *T. gondii* which through the innate immune sensing of vRNA, could drive a type I IFN-mediated inflammatory response. While we cannot establish this mechanism causally, it is plausible and supported by the body of available mechanistic and transcriptomic evidence.

A link between neurotropic toxoplasmosis and neuroinflammatory and neurodegenerative conditions, such as Alzheimer’s disease (AD) has been proposed (Nimgaonkar et al., 2016; Nayeri Chegeni et al., 2019; Tyebji et al., 2019; Yang et al., 2021). More recently, more mechanistic studies have been able to link CNS *Toxoplasma* infection with several correlates of AD pathology - disruption of the BBB, glial activation and synapse loss, *inter alia* (Li et al., 2019; Ortiz-Guerrero et al., 2020; Castaño Barrios et al., 2021; Carrillo et al., 2023; Anaya-Martínez et al., 2023). The picture however remains far from conclusive. Other epidemiological work have found no significant effect of *T. gondii* seropositivity on cognitive function in adult patient populations (Wyman et al., 2017; Torniainen-Holm et al., 2019). In pre-clinical models, evidence has been provided for neuroprotective effects in AD mouse models (Jung et al., 2012; Cabral et al., 2017), while others still find no significant effect on murine age-related cognitive decline (McGovern et al., 2020). This underscores the importance of a better understanding of factors which influence *Toxoplasma* pathogenicity, which could explain why some patients develop CNS pathology, while others are seemingly unaffected, and even protected. In AD, CNS inflammation is balanced between beneficial (e.g., priming microglia for amyloid plaque clearance) and detrimental (e.g., synaptic loss, immune infiltration) functions (Heneka et al., 2015; Leng and Edison, 2021). *Toxoplasma* strain-dependent effects may go a long way towards clearing up the somewhat mixed picture observed to date (Cabral et al., 2017; Xiao et al., 2022). The presence of an *Apocryptovirus*, or similar PPV, is well-placed to skew the inflammatory equilibrium in chronic toxoplasmosis towards neuropathology, particularly when one considers the recent identification of a detrimental role for type I IFN signalling in AD (Roy et al., 2020, 2022; Sanford et al., 2023).

The strength of computational virology is in its capacity for searching massive swathes of data and identify novel and unexpected relationships between viruses and diseases. As the number of newly discovered viruses continues to grow, analysis for developing specific and evidence-based hypotheses can help focus where resource intensive biological experiments should be allocated. Ao and neuroinflammation is such a case study, and serves to demonstrate the need for, and merit of, the comprehensive characterization of Earth’s RNA virome.

## Materials and methods

### Querying the BioSample database

In the initial screen for novel and highly divergent neurotropic viruses, we searched the BioSamples SQL table in the Serratus SQL database (https://github.com/ababaian/serratus/wiki/SQL-Schema) with the query SELECT * FROM tismap summary WHERE biosample tags LIKE ‘%neuron%’ AND scientific name = ‘Homo sapiens’ AND percent identity *<* 90 ORDER BY coverage desc; and identified SRA run SRR1205923 in BioProject PRJNA241125 through manual curation. BLASTp and BLASTx (Altschul et al., 1990) were performed with a query of the u150420 palmprint identified in three separate libraries to measure the percent amino acid identity to known viral proteins using the non-redundant proteins database (nr). Furthermore, BLASTn against the nr/nt database was performed (date accessed: 2023-06-09).

### Viral genome identification, assembly, and endogenous virus evaluation

The three libraries (SRR1205923, SRR1204654, SRR1205190) identified through a Serratus SQL search were *de novo* assembled using rnaSPAdes 3.15.5 (Bushmanova et al., 2019). rnaSPAdes was run with parameters ‘--rna --s1 -t 64’. Palmscan (version 2, --threads 64 --seqtype nt) (Babaian and Edgar, 2022)—an RNA-dependent RNA Polymerase detector—was used to identify the viral RdRp contig in the assembled transcript file. For each library, the contigs identified by Palmscan were re-analyzed through ORFfinder (NCBI RRID:SCR 016643) and the longest ORF was extracted. This amino acid sequence was identical across both mRNA libraries SRR1205923 and SRR1204654, but contig in SRR1205923 had a low-coverage insertion which introduced a frameshift mutation which was manually corrected to the consensus sequence to result in an identical coding sequence to the other libraries. To test if the ORF encoded a plausible RdRp, the structure was predicted with Colabfold (AlphaFold mmseq2, v1.5.5) (Mirdita et al., 2022), using the putative RdRp ORF from SRR1205923. BWA-mem 0.7.17 -t 64 (Li and Durbin, 2010) was used for aligning SRR1205923 transcripts generated by the assembler back to SRR1205923’s reads (FASTQ); alignment was visualized in IGV 2.16.0 (Robinson et al., 2011).

The *T. gondii* RUB and *T. gondii* COUGAR DNA-seq data was downloaded from their respective BioProjects (PRJNA61119 and PRJNA71479). Bowtie2 (version 2.5.1, ‘--local --very-sensitive-local --threads 64 -q -x -U’) (Langmead and Salzberg, 2012) was used to map the DNA-seq reads to a reference database of Ao contig 1 and 2, and no reads were alignable.

### Identification of Ao contig 2 via host read-depletion

Reference genomes for human(GRCh38, accessed from NCBI (https://www.ncbi.nlm.nih.gov/datasets/genome/GCF_000001405.26/) and *Toxoplasma gondii* ME49 (TgondiiME49, accessed from ToxoDB (Gajria et al., 2008)) were downloaded and concatenated into one reference dataset (hgTg). Assembled contigs (rnaSPAdes) for each of the 4 RUB mRNA Nĝo libraries (SRR1205923, SRR1204654, SRR1204653, SRR1204652) were aligned to hgTg with Bowtie2 (version 2.5.1, ‘--local --very-sensitive-local --threads 64 -f -x -U’). SAMtools 1.17 view with parameters ‘-S -b -@64’ converted the SAM to a BAM filZe and another SAMtools view command with parameters ‘-f 4 -b -@ 64’ removed mapped contigs, retaining non-human, non-toxoplasma mapping contigs. The 4 libraries SRR1205923, SRR1204654, SRR1204653 and SRR1204652 had 76, 92, 52 and 50 unmapped contigs respectively. All unmapped contigs were converted to FASTA with SAMtools and were then aligned against NCBI’s nr proteins database using DIAMOND 2.1.8 (Buchfink et al., 2015) in BLASTx mode with parameters ‘--masking 0 --unal 1 --mid-sensitive -l 1 -p14 -k1 –threads 64 -f 6 qseqid qstart qend qlen qstrand sseqid sstart send slen pident evalue full qseq staxids sscinames’. The contigs with the highest coverage matched *E. coli* sequence, the 3rd highest by coverage was the Ao RdRp contig and the 2nd highest across all 4 RUB libraries was a *≈* 1280 nucleotide long contig with no known associations through BLAST. This unknown contig was common throughout all 4 RUB libraries and was called the putative contig 2. Subsequently we show this contig correlated with contig 1 (RdRp) across the Ngo BioProject, and we identified co-evolving contig 2 homologs in 19 related *Apocryptovirus sp.*, supporting that contig 2 is viral in origin.. Subsequently, a genome map representation was created for both contig 1 and contig 2 with gggenes 0.4.1 (Wilkins, 2020), an R 4.2.2 package for drawing gene arrow maps and ggplot2 3.4.0 (Wickham, 2016). Finally, a Pearson correlation coefficient was calculated with ggpubr 0.6.0 in R (Kassambara, 2023).

### Assembling and quantifying of Ngô et al. Libraries

117 out of the 237 libraries in the Ngô et al. 2017 BioProject are labelled as mRNAseq data, the other 120 libraries are micro RNA (miRNA). The 117 mRNAseq libraries were pre-fetched and the FASTQ files were extracted using SRAtoolkit 3.0.5. Next, all 117 Ngô mRNA libraries were *de novo* assembled with rnaSPAdes with parameters ‘--rna --s1 -t 64’. Using Bowtie2 ‘-build’ a dataset was created with all 4 genomes/sequences: Human (HgCh38), *T. gondii* ME49 (Tg64), Ao contig 1 RdRp, Ao contig 2. Next, the reads for all 237 libraries in Ngô *et al*. were aligned against this dataset using Bowtie2 with parameters ‘—local --very-sensitive-local --threads 64 -q -x -U’. To convert the SAM file to a BAM file, SAMtools with parameters ‘-S -b -@ 64 -F 260’ were used. Flag -F 260 removes all unmapped reads and non-primary alignments. The output of SAMtools was then piped into Seqkit 2.4.0 (Shen et al., 2016) with the ‘-c’ parameter which counts the mapped reads per chromosome/sequence.

### Assembling and quantifying Melo et al. Libraries

Similarly to the approach taken with libraries from Ngô *et al*., the FASTQ files for all 32 libraries were downloaded using SRAtoolkit for the BioProject from Melo *et al*. 2013. All 32 Melo libraries were de novo assembled with rnaSPAdes parameters ‘--rna --pe -t 64’. Bowtie2 was used to create a dataset with all 4 genomes/sequences: Murine (GRCm39), Tg64, Ao contig 1 RdRp, Ao contig 2. Next, the reads for all 32 libraries in Melo *et al*. were aligned against this dataset using Bowtie2. To convert the SAM file to a BAM file, SAMtools was used (see: *Assembling and quantifying of Nĝo et al. Libraries* above for parameters). All the metadata for the entire Melo *et al*. BioProject (PRJNA241125) was collected with Google’s BigQuery tool. Using this metadata and the data parsed with Pandas 2.0.3 (pandas development team, 2020), Figure 3 was generated in R with dplyr and ggplot2.

### Screening for and assembling the Apocryptoviruses

To identify RdRp related to Ao, we aligned the sOTU in palmDB (version 2023-04) Babaian and Edgar (2022) to the Ao palmprint (usearch -usearch global palmDB.palmprint.faa -db ao.palmprint.faa -id 0.3), yielding matches in ‘u380516’, ‘u476932’, ‘u706419’, ‘u492272’, ‘u602981’, ‘u1051092’, ‘u845773’, ‘u857620’, ‘u665910’, ‘u819619’, ‘u584295’, ‘u691934’, ‘u71279’, ‘u145522’, ‘u626963’, ‘u150420’, ‘u592253’, ‘u828652’, ‘u617666’, ‘u964003’, ‘u643849’, ‘u993146’, ‘u770301’, ‘u533578’, ‘u460145’, ‘u419915’, ‘u849389’, ‘u1004674’, ‘u942391’, ‘u10732’, ‘u761722’, ‘u599206’ and ‘u1009116’. To query which Serratus-processed SRA runs contained microcontigs matching these palmprints, we performed the SQL query ‘SELECT * FROM palm sra2 WHERE qc pass = ‘true’ AND sotu IN (‘u380516’, ‘u476932’, ‘u706419’, ‘u492272’, ‘u602981’, ‘u1051092’, ‘u845773’, ‘u857620’, ‘u665910’, ‘u819619’, ‘u584295’, ‘u691934’, ‘u71279’, ‘u145522’, ‘u626963’, ‘u150420’, ‘u592253’, ‘u828652’, ‘u617666’, ‘u964003’, ‘u643849’, ‘u993146’, ‘u770301’, ‘u533578’, ‘u460145’, ‘u419915’, ‘u849389’, ‘u1004674’, ‘u942391’, ‘u10732’, ‘u761722’, ‘u599206’, ‘u1009116’)’, returning the 185 matching libraries (Supplementary Table S3). Sorting by microcontig coverage for each palmprint-library, FASTQ files from the top three libraries were downloaded using SRAtoolkit and assembled using rnaSPAdes with parameters ‘--rna -t 64 --pe’. RdRp-contigs were identified as described above in each library.

### MSA and phylogenetic tree of Ao and related Narnaviruses

In order to sample Ao-related viruses, we first searched for available sequences in GenBank’s non-redundant protein (nr) database, using PSI-BLAST and default parameters with the BLOSUM62 substitution matrix (date accessed: June 2023). We queried the database with the entire amino acid sequence for the Ao RdRp recovered from the RUB strain of *T. gondii* (SRR1205923); all resulting sequences had E-values *<* 7*e −* 28. Notably, this approach failed to capture more distantly related Narnaviruses, such as those previously described in arthropod metagenomic samples, yeast, and nematodes. Thus, we supplemented the MSA with RdRps from an additional BLASTp search on *Coquillettidia venezuelensis* Narnavirus 1 (QBA55488.1). For a more distant clade, we selected 7 ICTV-recognized *Mitoviridae* species, two from each genus except *Kvaramitovirus*, for which there is only one accepted species. As an outgroup, we added two species from the family *Fiersviridae*: bacteriophages MS2 and Qbeta. The sequence with the highest percent identity relative to Ao RUB among these was QIM73983.1 at 53.53% identity, but only 40% coverage. For this reason, after aligning the sequence with MUSCLE v3.8.1551 (Edgar, 2004), we decided to manually trim the MSA to the RdRp palm (motifs F-E) and thumb subdomains. This ensured partial contigs could be fairly represented in the phylogeny. Some BLAST hits, however, were too incomplete to satisfy this requirement and were therefore removed.

With the closest relative in GenBank at such a low identity, we re-queried the SRA for Ao-like sequences. Using DIAMOND, a sequence database of all Ao-like palmprint sOTUs was created. We then aligned all assembled transcripts from Ao-like RdRp-containing libraries to the Ao-like palmprints (DIAMOND, ‘-blastx --very-sensitive --threads 64 -f 6 qseqid sseqid pident length mismatch gapopen qstart qend sstart send evalue bitscore full qseq’). The alignment outputs were merged and transcripts with 100% identity to the palmprint, *≈* 3000 nt length, and the highest coverage if multiple choices were available, were chosen as the representative contigs for each virus. 8 of the 55 libraries did not have a high quality representative contig, and were discarded. The remaining contigs were analyzed with ORFfinder command line to find the ORFs. The longest ORF was chosen, and then confirmed to be the RdRp using BLASTp. These RdRp ORFs were combined into one FASTA file, and are the source of Ao-like virus RdRp. The final MSA contains 169 RdRps; a full list of the selected sequences, their GenBank/SRA accessions, and associated metadata is available in the Supplementary Materials. Based on the RdRp palm, we generated a maximum-likelihood phylogenetic tree using IQ-TREE v2.2.2.7 (Minh et al., 2020) and viewed in ggtree v3.10.0 (Yu et al., 2016), where the substitution matrix was selected as rtREV+F+I+R8 by ModelFinder (Kalyaanamoorthy et al., 2017). Bootstrap values on 1000 replicates were generated via UFBoot v1 (Hoang et al., 2018).

### Novel Ao-proximal RdRp Motif Identification

Based on the phylogeny, sequences displaying high-relatedness with Ao were selected. The full sequences (as opposed to just the palm and thumb) were retrieved and aligned with MUSCLE. A conservation map showing alignments for selected sequences in the MSA was generated using ggmsa v1.8.0 (Zhou et al., 2022a). Subsequences with high occupancy and conservation were selected as novel motifs *α*-*µ*. We also selected “non-motif” subsequences to serve as a background. These constituted the high occupancy regions between motifs. Sequence logos for the novel and canonical RdRp motifs were generated via WebLogo v2.8.2 (Crooks et al., 2004). Pairwise conservation was calculated based on the percentage of residues with similar chemical properties between two aligned subsequences. No penalties were given to mismatched residues or gaps. A matrix of pairwise comparisons between all motifs across all selected sequences was calculated with Pandas. All FASTA files and the Jupyter notebook used for calculation are available in the supplementary data. Mean conservation across the novel, core, and non-motifs was calculated and used as the basis for a scatterplot in R using ggplot2. Deltas (differences between mean values for the novel, core, and non-motif sets) were also plotted with ggplot2 in the form of a violin plot.

### RdRp structure analysis

Protein structures of full-length RdRps were predicted using Colabfold for the Apocryptoviruses and select Apocrypto-proximal RdRps. Structures were visualized in PyMol (Schrodinger, LLC, 2015). The topology map was generated with PDBSum (April 10, 2023 version) (Laskowski et al., 2018) and further edited with Adobe Illustrator. TM-align v20170708 (Zhang and Skolnick, 2005) was used for structural alignments, and the ColorByRMSD plugin (April 6, 2016 version) for PyMol was used to generate the distance maps.

### Contig 2 recovery

To recover contig 2 for each Ao-like virus, a DIAMOND database of all 4 mRNA RUB Nĝo Ao contig 2’s was created. Using BLASTx, the transcripts for all Ao-like viruses were aligned against this database. Pandas was used to sort and combine all the alignment data. For each library, the contig with the best alignment identity, e-value, coverage, and length was chosen. These contigs were combined into one FASTA file. This combined file was then put through the ORFfinder command line and all the ORFs were extracted. An MSA of the extracted ORFs was created using MUSCLE. Upon viewing the alignment, 2 profiles were generated: one for contig 2’s pORF1 and the other for ORF2. Based on these profiles, 2 HMM models (one for ORF 1 and one for ORF 2) were created with HMMER3 v3.4 Using these HMM models, HMMER’s hmmsearch was used to search all Ao-like virus libraries. Any ORFs that hit against the HMM model were labelled either pORF1 or pORF2 according to which HMM model they aligned against.

### Differential gene expression

To begin, the human reference genome (GRCh38.p14), its corresponding gene annotation, as well as the FASTQ files of the Ngô dataset accessions for Macrophage (MonoMac6) and Neuronal Stem Cell (NSC) were downloaded. Initially, all experiments were performed on macrophages, and then replicated with the NSC dataset. HISAT2 v.2.21 (Kim et al., 2019) was used to extract splice sites and exons, and then used to index the human genome to generate SAM files for each accession. SAM files were converted to BAM files using SAMtools (See Materials and Methods, *Assembling and quantifying of Nĝo et al. Libraries*). A matrix of counts per human gene symbol was generated using the featureCounts tool in the SubRead v.2.14.2 package (Liao et al., 2014). DESeq2 v.1.40.2 (Love et al., 2014) was used to analyze the count matrix and test for differential gene expression. DESeq2 and ggplot2 were used to create MA plots, PCA plots, and volcano plots with highlighted values having Benjamini-Hochberg adjusted p-value *<* 0.05. Upon further analysis, it was noted that of the four control mock strains used, two appeared to be pro-inflammatory (Supplementary Figure S8). To investigate this, a spreadsheet consisting of all of the metadata from the Ngô *et al*. experiment was created. From this, it was observed that Ngô *et al*. performed their experiments in separate runs. This batch effect explains the variation in the immune system response for the control Mock strains. To account for this batch effect, any samples that were used in batches that presented hyper-inflammatory symptoms were not used for further analysis. To determine whether the defined set of genes showed statistically significant differences between phenotypes, we used the Gene Set Enrichment Analysis (GSEA v.4.3.2) software (Subramanian et al., 2005). Using the Molecular Signatures Database (MsigDB) (Liberzon et al., 2015), we focused on the hallmark collection of gene sets, as well as five non-hallmark gene sets. The five non-hallmark gene sets were chosen on the basis of containing the genes *IFNB1*, *IFNW1*, *IFNA13*, *IFNA1*, *IFNE*, *IFNL3*, *IFNL4*, all of which were the interferon genes being upregulated amongst *T.gondii* strain infection. Using all of these gene sets, GSEA data matrices were then used to create both enrichment plots and heat maps of NES scores.

### Quantifying co-occurrence of Apicomplexa presence with Apocryptoviruses

We selected the six, non-laboratory source organisms in which apocryptoviruses were identified. Each of these sets was further partitioned into a contigency table of four groups according to two categorical variables. The first variable was whether or not there is significant Apicomplexa signal (*>* 128 matching reads) within that SRA run, as reported by STAT under the phylum *Apicomplexa* (BigQuery: SELECT * FROM nih-sra-datastore.sra tax analysis tool.tax analysis WHERE tax id = 5794; Data was retrieved on July, 2023.). The second variable was whether or not the SRA run belongs to the proposed list of identified Apocryptoviruses (Supplementary Table S3). To test if these variables were associated, we performed a Fisher’s Exact Test (scipy.stats.fisher exact), results are summarized in Figure 5.

### Reverse Transcriptase PCR for A. odysseus in cultured T. gondii

Human foreskin fibroblasts (HFFs) were cultured in Dulbecco’s Modified Eagle Medium from (Invitrogen, cat# 11965-118) supplemented with 10% heat-inactivated fetal bovine serum (FBS, Genesee Scientific, cat# 25-550), 2 mM L-glutamine, and 50 µg/ml of both penicillin and streptomycin, along with 20 µg/ml gentamicin. *Toxoplasma* strains RH (Ao-control), RUB (Ao+), and COUGAR (Ao+) from (Melo et al., 2013) were propagated in HFFs in T25 culture flasks. RNA was extracted using the RNeasy Plus Mini Kit (Qiagen, cat# 74134), following the manufacturer’s instructions. The yield and purity of the extracted RNA were assessed using a Nanodrop. cDNA was synthesized using the Reliance Select cDNA Synthesis Kit (Bio-Rad, cat#12012802) for 20 minutes at 50°C. Amplification of *A. odysseus* RUB/COUGAR was performed using primers: Forward Primer - 5’-ATTGTTCCCGTGCATGACTG-3’ and Reverse Primer - 5’-TCTTGAGAGTCCGGCTTTGG -3’. As a loading and positive control for cDNA quality, *Toxoplasma GRA1* amplification was performed with primers: Forward Primer - 5’-TATTGTCGGAGCTGCTGCATCG-3’ and Reverse Primer - 5’-GCTCACTGCATCTTCCAGTTGC-3’. To test if Ao is a potential endogenous viral element, the same reaction without reverse transcriptase was performed for all samples. PCR reactions with cDNA from RH, RUB, and COUGAR, and a no-template negative control were also included. The PCR was performed using MangoMix with MangoTaq (Bioline, cat# C755G90), for 34 cycles of; denature: 15 seconds at 95°C, anneal: 30 seconds at 61°C (*A. odysseus*) or 59°C (GRA1), and extension: 30 seconds at 72°C. Amplicons were purified using the DNA Clean and Concentrator kit (Zymo Research,cat# 11-305C) and sequenced by QuintaraBio (3563 Investment Blvd, Suite 2, Hayward, CA 94545) with aforementioned primers. The *A. odysseus* RUB and COUG sequences were a 100% match to A.odysseus SRR446933 and A.odysseus SRR446909, respectively.

## Acknowledgments

We would like to thank R. Valencia, J. Shen, J. Parkinson, J. Charon, and A. Reinke for their helpful comments on the manuscript. This work is supported by a Project Grant from the Canadian Institutes for Health Research (CIHR PJT - 190150). AH is supported by the University of Toronto’s Department of Molecular Genetics undergraduate research opportunity program (UROP). JC is supported by the University of Toronto’s Department of Molecular Genetics undergraduate research opportunity program (UROP). We are grateful to the entire team managing the NCBI SRA and the biology community for openly sharing their data, with special thanks to the McLeod lab for the RUB Toxoplasma data. Computing resources were provided by the University of British Columbia Community Health and Wellbeing Cloud Innovation Centre, powered by AWS.

## Data availability and supplementary data

Apocryptovirus sequences are available in GenBank under BioProject PRJEB71349. All sequences and supporting data are available at https://github.com/ababaian/serratus/wiki/Apocryptovirus. Project notebooks and code are available via S3 (https://mtnemo.s3.amazonaws.com/README.md).

## Supplementary Tables

**Supplementary Table S1. Ngô sequencing datasets and meta-data.** Metadata describing BioProject PRJNA241125 (Ngô study) including flowcell header metadata extracted from raw (fastq) sequence headers and normalized expression (RPKM) of *Apocryptovirus odysseus* per library.

**Supplementary Table S2. *Toxoplasma gondii* strains.** Strains of *Toxoplasma gondii* analyzed from BioProjects PRJNA241125 (Ngô study) and PRJNA114693 (Melo study), and their associated meta-data.

**Supplementary Table S3. Apocryptoviruses and associated SRA metadata. A.** Complete list of exemplar *Apocryptoviruses* identified in this study. **B.** Description of each Sequence Read Archive (SRA) library from which a relevant species-like operational taxonomic unit (sOTU) was identified. **C.** The pairwise percentage identity between exemplar *Apocryptoviruses*. **D.** Duplicate of Table 1 in spreadsheet format

**Supplementary Figure S1.**
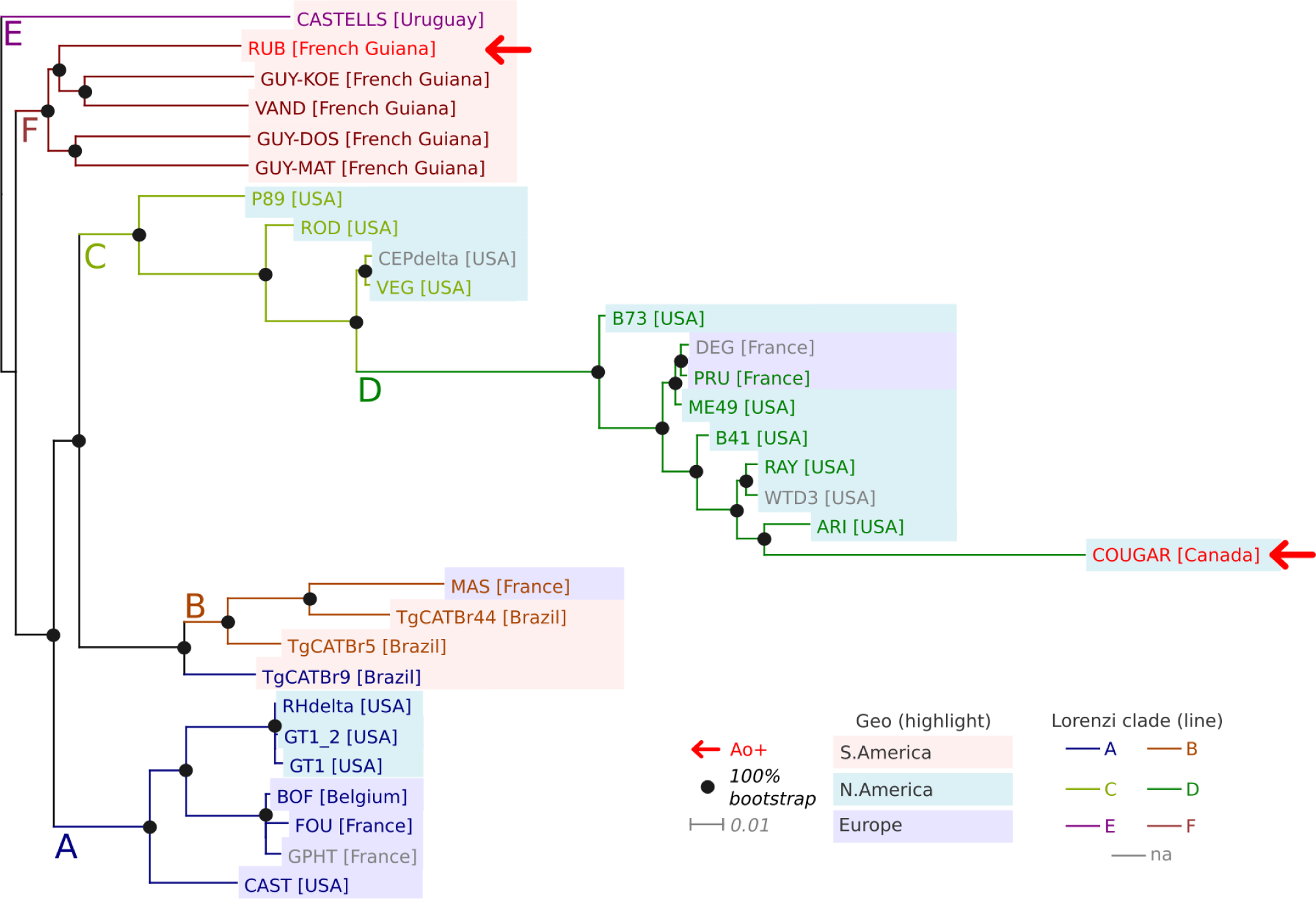
Maximum likelihood tree for T. gondii transcriptomes. RNAseq reads were aligned to the T. gondii ME49 reference genome (version 64) from each of the 31 T. gondii samples of the Melo study (Melo et al., 2013). Phylogeny was constructed from consensus single nucleotide polymorphisms (SNPs) in expressed annotated exons. 8,201,735/30,124,248 (27.22%) exonic sites were expressed across all samples of which a further 156,773/8,201,735 (1.91%) sites were polymorphic in at least one sample. IQ-TREE (v2.2.2.6; iqtree2 -s melo.snp-only nogap msa.fa -m TEST+ASC -bb 1000 -alrt 1000; model: ‘TVM+F+ASC+G4) was ran. Nodes supported by 100% bootstraps are indicated with a black circle.

**Supplementary Figure S2.**
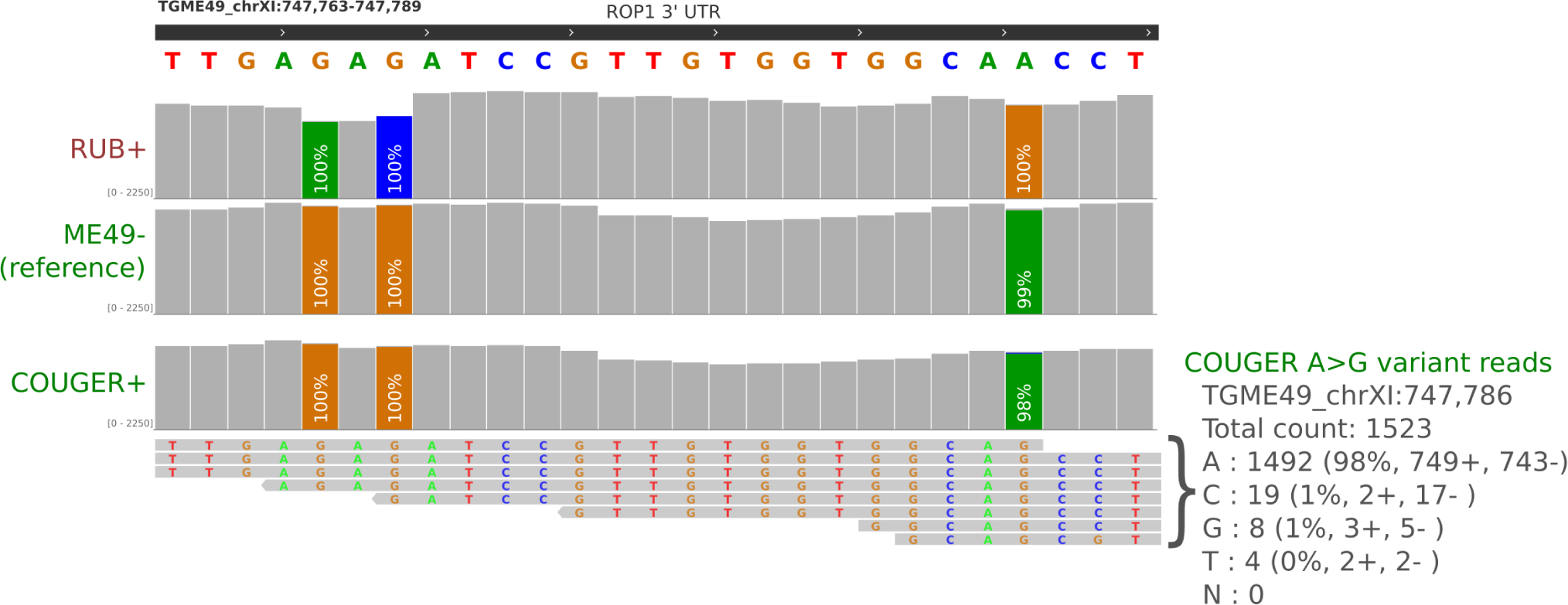
Melo study T. gondii transcriptomes sample contamination. RNAseq reads were aligned to the T. gondii ME49 reference genome (version 64) from each of the 31 T. gondii samples of the Melo study (Melo et al., 2013). To test if the presence of Ao in RUB and COUGAR was the result of sample cross-contamination, we manually inspected reads from the Melo transcriptomes in IGV 2.16.0. The ME49 strain is included as an Ao-negative control related to COUGAR (Lorenzi clade D). RUB-specific alleles were not present above the sequencing-error background (°1 %) in the COUGER sample or vice versa. Where possible, the reads for potential cross-contaminating alleles were inspected at adjacent linked sites. Manual inspection of several dozen sites revealed no read-level evidence for cross-contamination of samples.

**Supplementary Figure S3.**
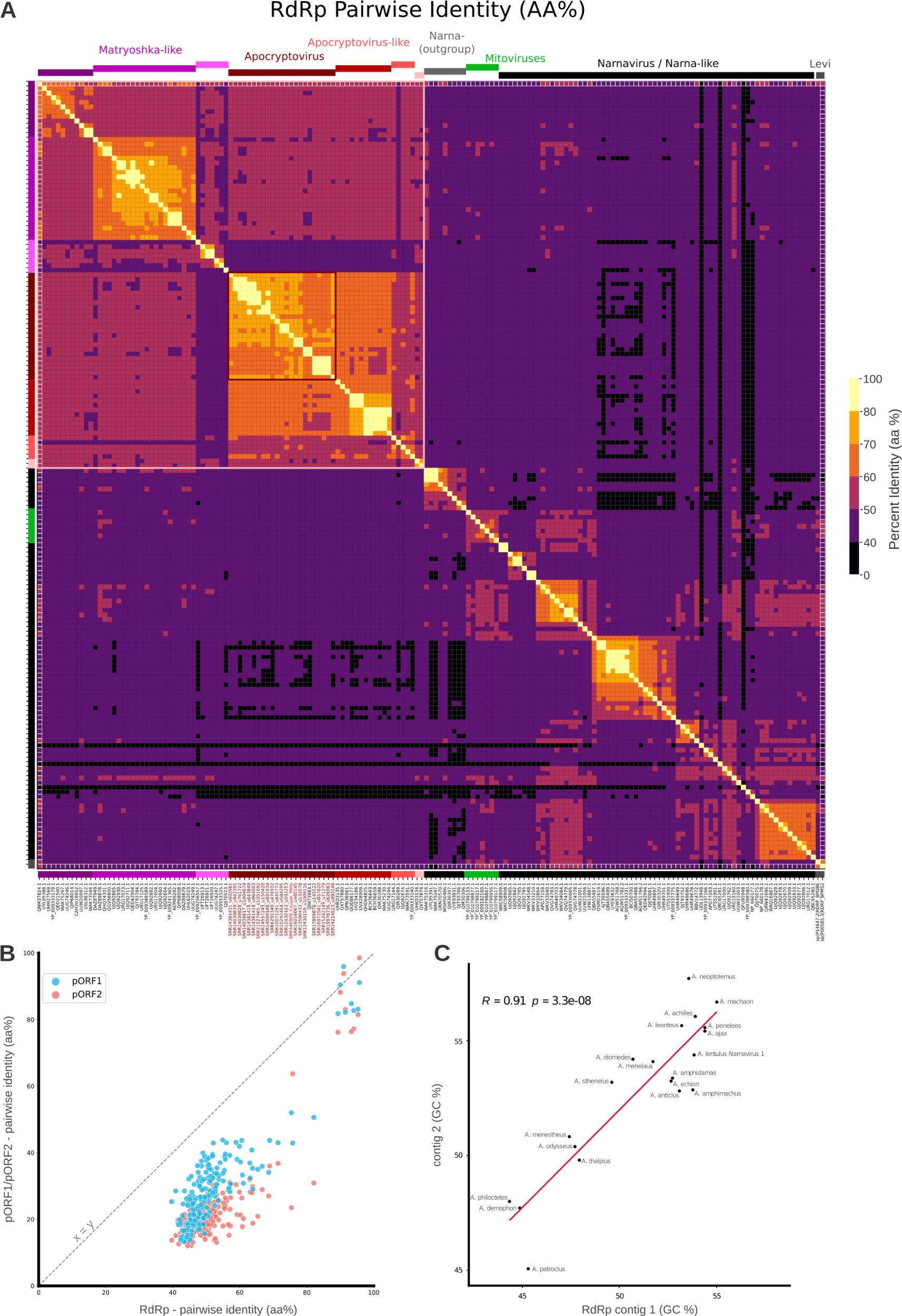
Pairwise percent amino acid identity between Narnavirus RdRp A. Heatmap plotting the pairwise percent identity for each RdRp in the narnavirus phylogeny. Percent identity was calculated on the basis of pairwise alignments for the RdRp core (motifs F-E) and thumb, gaps were not penalized. The heatmap is symmetrical (i.e. RdRps on the diagonal are identical), with tentative phylogenetic groupings shown. **B.** Scatterplot of between-*Apocryptovirus* (pairwise) percent identity for RDRP, pORF1 or pORF2 where contig 1 and contig 2 association was unambiguous. USEARCH v11 was run with parameters ‘-allpairs global -acceptall’. Scores were extracted using Pandas and plotted with Matplotlib v3.8 (Hunter, 2007). **C.** Pearson correlation of GC content across contigs 1 and 2 for each Apocryptovirus species. *A. lentulus* Narnavirus 1 is also included as a closely-related bi-segmented narnavirus.

**Supplementary Figure S4.**
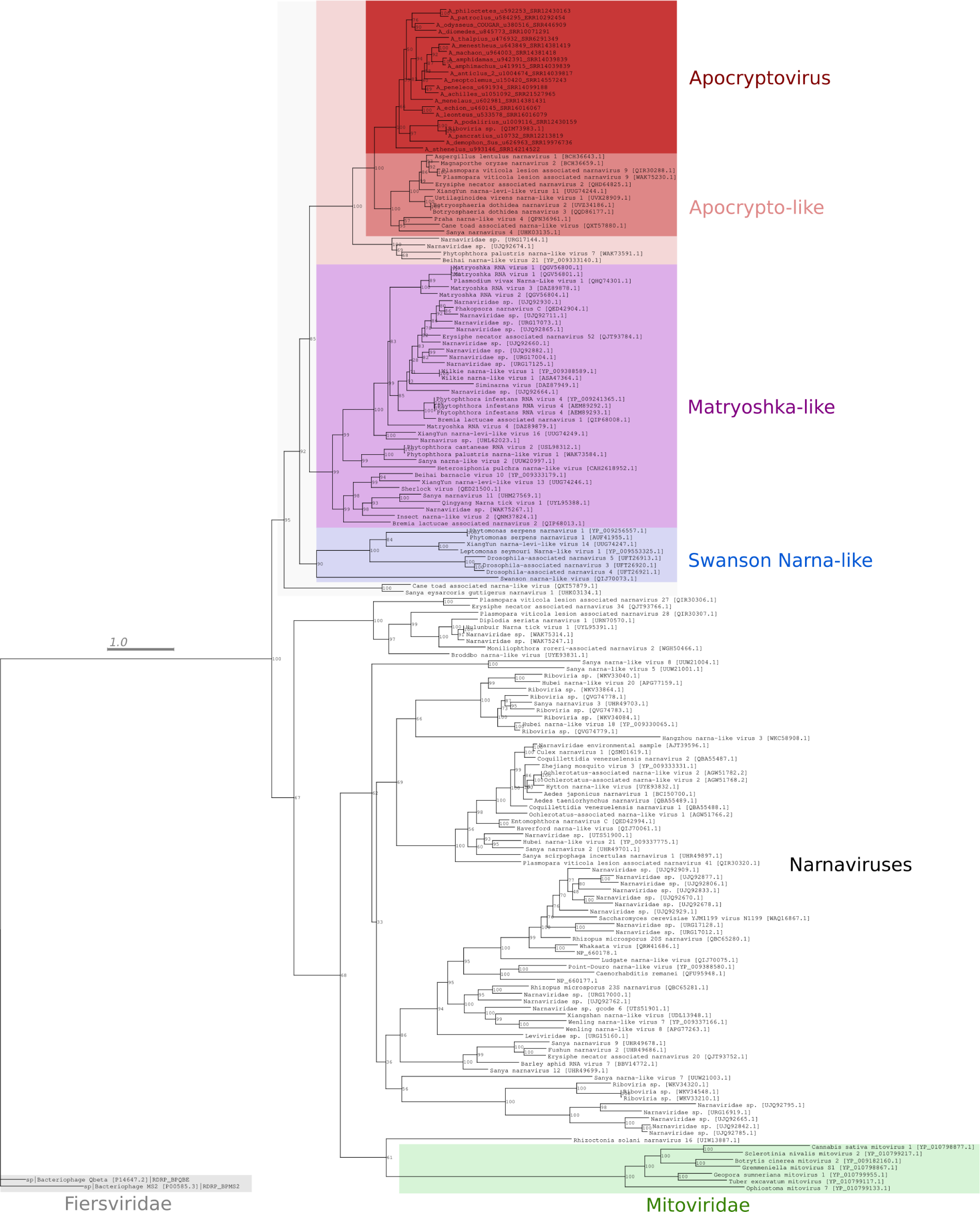
Narnavirus RdRp phylogeny with Apocrytoviruses. Maximum-likelihood phylogenetic tree of *A. odysseus* and the related *Apocryptoviruses* estimated based on RdRp palm (motif F-E) and thumb subdomains. *Apocryptovirus* is placed with high confidence (100% bootstrap support) in a clade of *Apocrypto*-like Narnaviruses, which include the bi-segmented Aspergillus lentulus narnavirus 1 (BCH36643.1), which itself are a sister-clade to the Matroyshka and Matryoshka-like Narnaviruses. Bacteriophage (Fiersviridae) is included as an out-group. Scale bar represents 1 amino acid substitution per site.

**Supplementary Figure S5.**
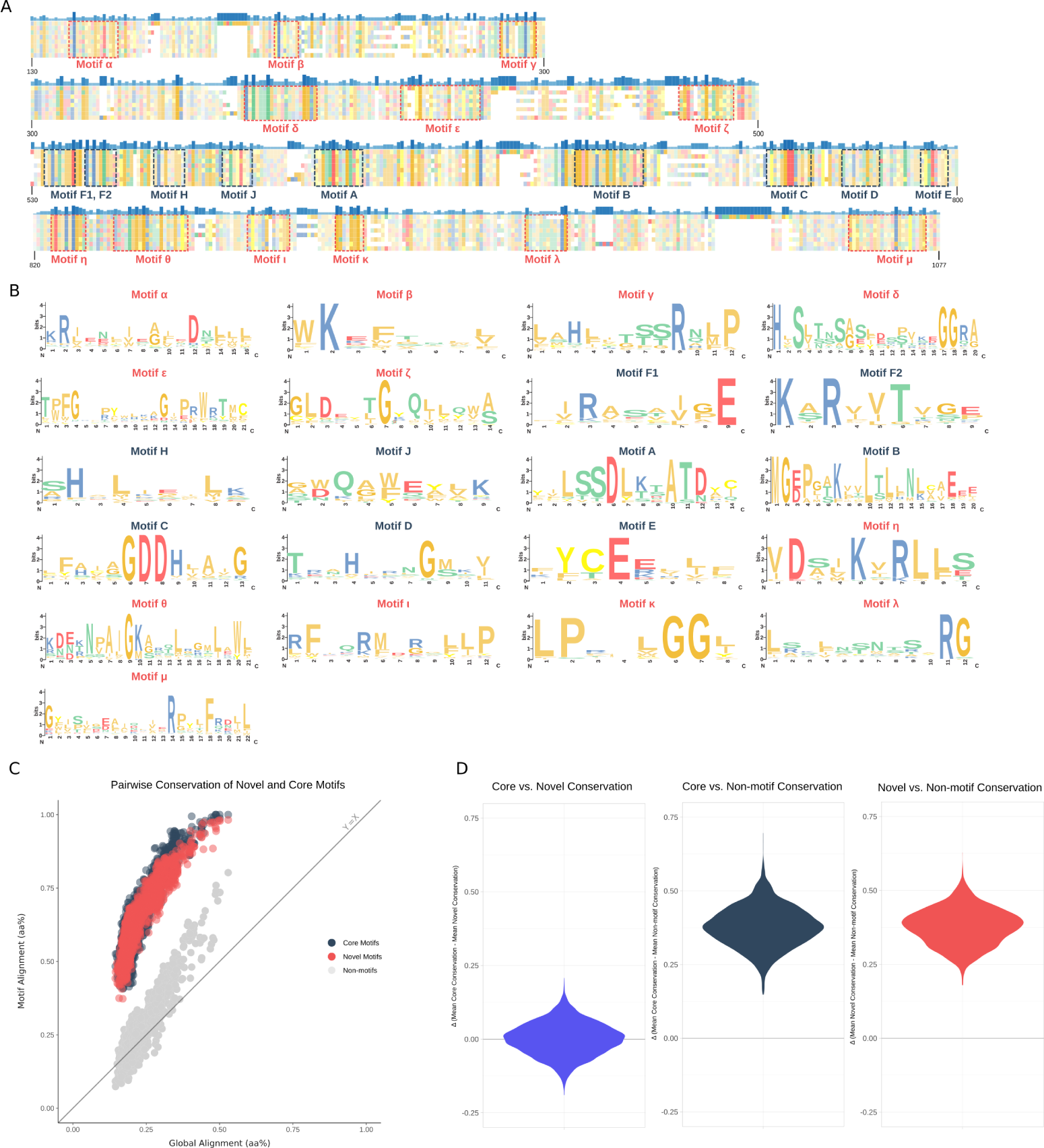
Novel RdRp Motif Sequence Analysis A. Conservation map of selected RdRps from “Ao-proximal” RdRp MSA. Sequences as follows (from top to bottom): Broddbo narna-like virus [UYE93831.1], Aspergillus lentulus Narnavirus 1 [BCH36645.1], Matryoshka RNA virus 3 [DAZ89878.1], Ao [SRR446909], Matryoshka RNA virus 4 [DAZ89879.1], Ao-like sp. [SRR14557243], Cane toad associated narna-like virus, [QXT57879.1], Beihai narna-like virus 21 [YP 009333140.1]. Novel and core motifs are highlighted, residues colored by chemical properties and degree of conservation. **B.** Sequence logos for novel and core motifs. **C.** Scatterplot of mean pairwise percent conservation between novel and core motifs. Pairwise percent conservation was calculated within each motif for every sequence selected. The mean conservation was calculated across the core motifs (F-E) and the novel motifs (*α*-*µ*) and plotted for each pairwise comparison. Inter-motif regions with high occupancy are also plotted in the same manner, serving as a background conservation rate. **D.** Violin plots for the difference in mean conservation for groups specified. Strongly positive/negative values indicate a large difference in conservation. Values close to zero suggest both sets of motifs are equivalently conserved.

**Supplementary Figure S6.**
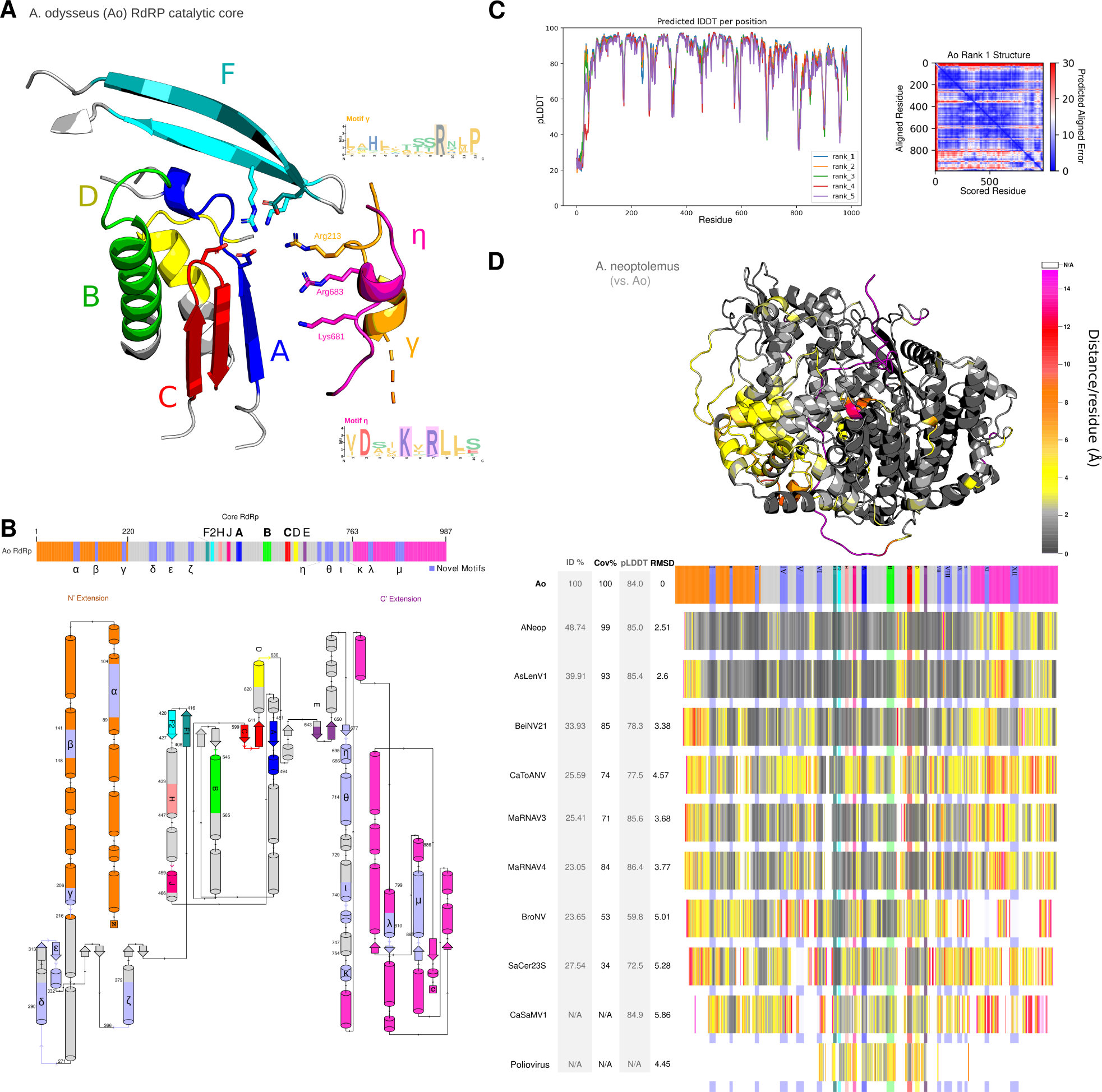
Extended Structural Analysis A. Predicted structure of Ao RUB RdRp. Investigating the ColabFold predictions, we note the positioning of motifs *γ* and *η* are in close proximity with the catalytic core of RdRp, with residues Arg9 of motif *γ*, together with Lys5 and Arg7 of motif *η*, are highly conserved across the Apocrypto viruses. Structurally, the three are oriented towards the catalytic site above motif C. The structure and evolutionary conservation suggests these might act as a stabilizing mechanism for the negatively-charged template RNA backbone. **B.** Secondary structure topology map for Ao core RdRp and extension domains with canonical and novel motifs highlighted. **C.** Plots for the predicted local difference distance test (pLDDT), a per-residue estimate of accuracy, for the five predicted structures and predicted alignment error for the rank one model. **D.** Distance maps for pairwise alignments of select RdRp structures against Ao. The structure for *A. Neoptolemus* is shown, with each residue colored by its distance to the corresponding residue in the structural alignment against Ao RdRp. Residues that did not align are shown in white. The N’- and C’-extensions appear to be topologically unique domains, distinct from the core RdRp. The motifs identified in these regions are structurally conserved within the Apocrypto-proximal RdRps, with the N’-motifs (*α*-*γ*) more strongly conserved than the C’-ones (*λ* and *µ*). Moreover, some of these (e.g., motif *β*) may be conserved across all of *Narnaviridae* and *Mitoviridae*, but it is difficult to say for certain as structural alignments may be capturing similarity, and not homology at these evolutionary distances. **E,F.** ColabFold prediction pLDDT scores for Ao RUB pORF1 homodimer and pORF2, respectively.

**Supplementary Figure S7.**
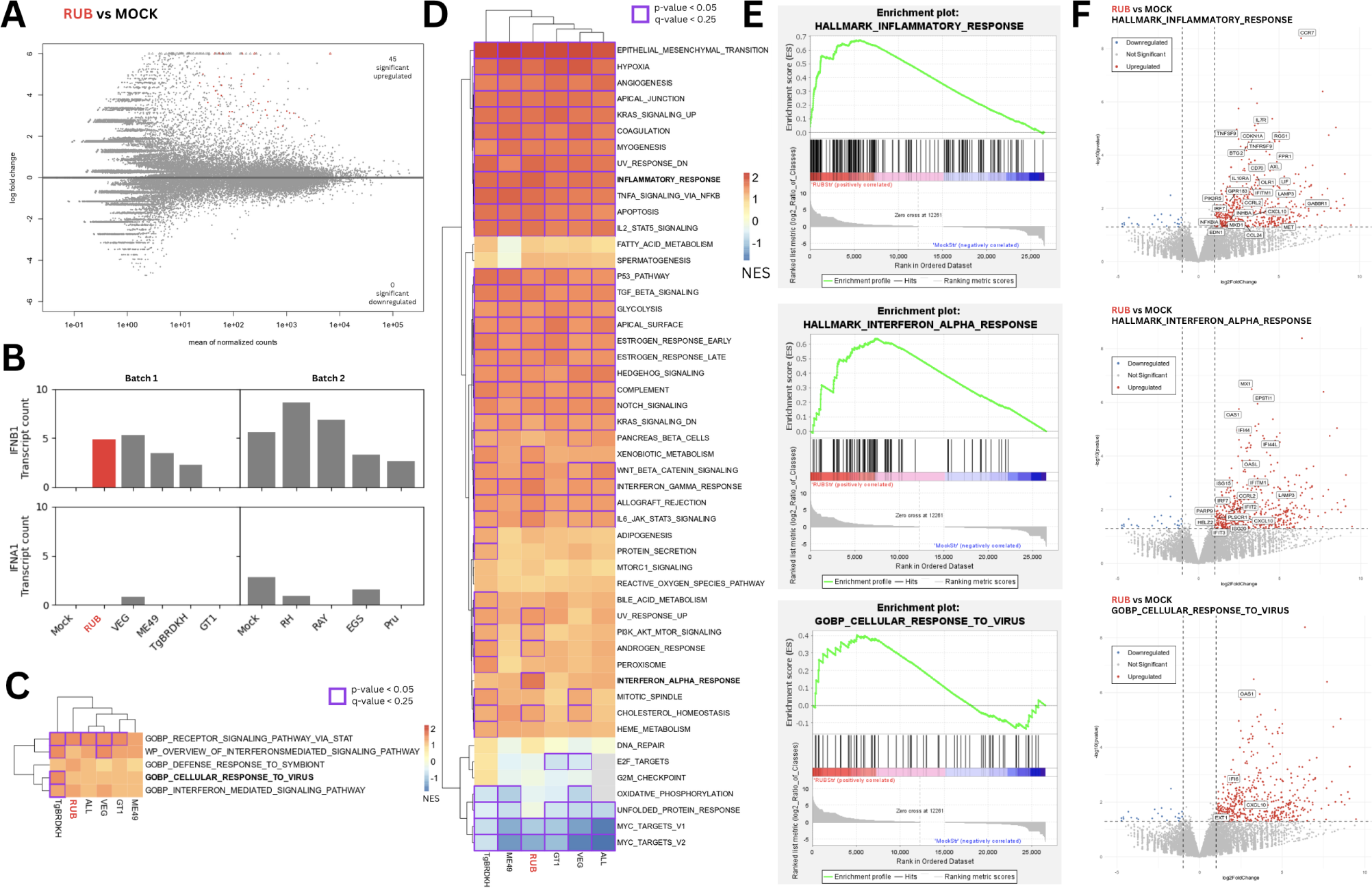
DGE of human macrophages infected with various *T. gondii* strains. **A.** MA plot of *T. gondii* - RUB vs mock genes (highlighted: Benjamini-Hochberg adjusted p-value *<* 0.05) **B.** Bar plot of normalized transcription counts of *IFNB1* and *IFNA1* across *T. gondii* strains and mock sequenced in Nĝo. et al experiments, separated by batch. **C.** Heat map of Normalized Enrichment Scores (NES) from Gene Set Enrichment Analysis (GSEA) using gene sets possessing interferon-specific genes, namely *IFNA1* and *IFNB1*, applied to the *T. gondii* strains. **D.** Heat map of NES values from GSEA using the Hallmark gene sets. **E.** GSEA curves comparing RUB vs. Mock strain using inflammatory response, cellular response to virus, and interferon-mediated signalling pathway gene sets. **F.** Volcano plot of differentially regulated genes with genes of notable gene sets being labeled.

**Supplementary Figure S8.**
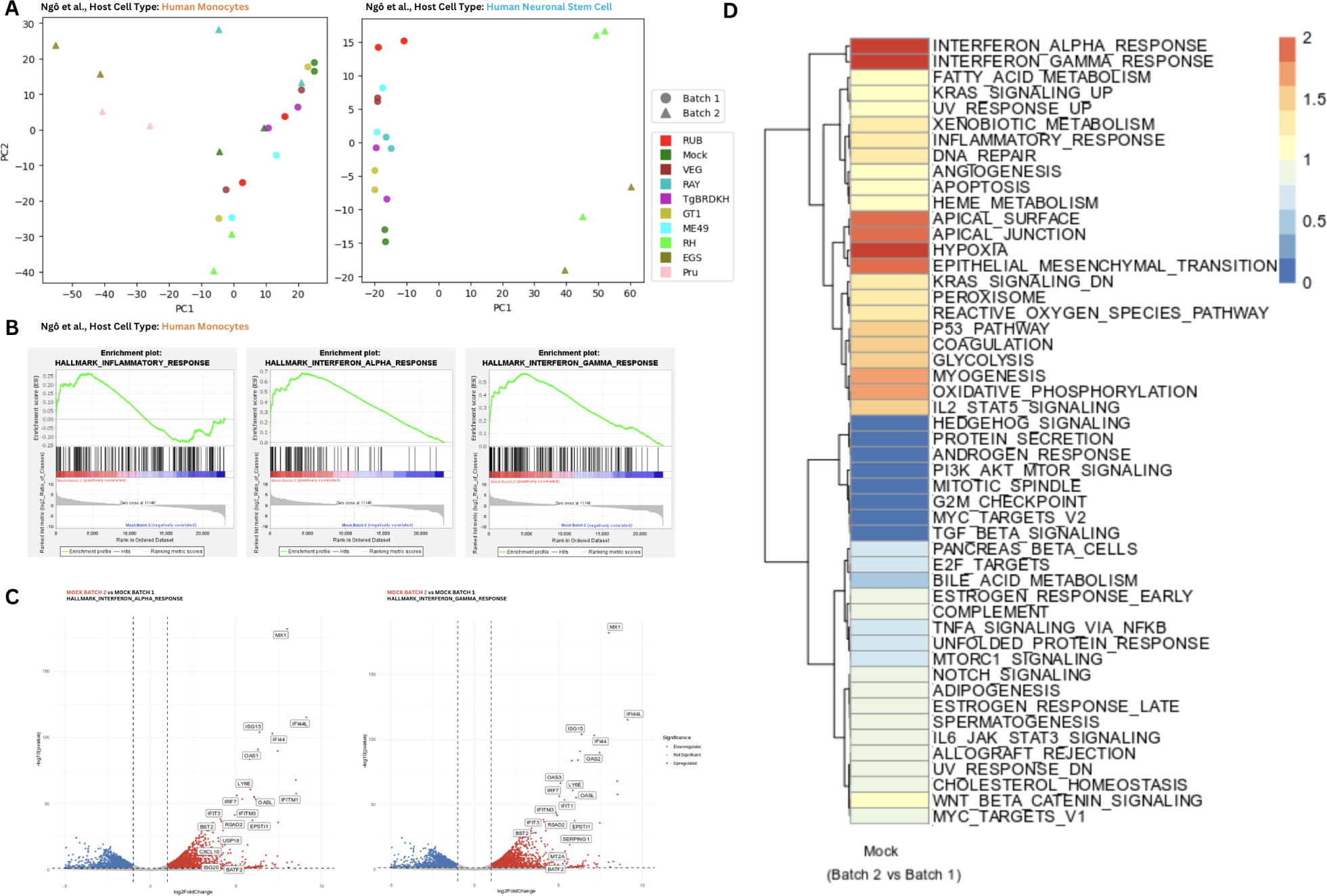
Inspection of Differential Gene Expression data displaying batch effects. **A.** PCA plot of batch effects visible in Nĝo *et al*. datasets for Human Monocytes and Neuronal Stem Cells. **B.** GSEA curves comparing Mock strains between Batches using inflammatory response, interferon alpha response, and interferon gamma response gene sets. **C.** Volcano plot of differentially regulated genes with genes of interferon alpha response, and interferon gamma response gene sets being labeled. **D.** Heat map of NES values from GSEA using the Hallmark gene sets comparing mock batch 1 vs. mock batch 2.

